# The Filter Detection Task for measurement of breathing-related interoception and metacognition

**DOI:** 10.1101/2020.06.29.176941

**Authors:** Olivia K. Harrison, Sarah N. Garfinkel, Lucy Marlow, Sarah Finnegan, Stephanie Marino, Laura Nanz, Micah Allen, Johanna Finnemann, Laura Keur-Huizinga, Samuel J. Harrison, Klaas E. Stephan, Kyle T. S. Pattinson, Stephen M. Fleming

## Abstract

The study of the brain’s processing of sensory inputs from within the body (‘interoception’) has been gaining rapid popularity in neuroscience, where interoceptive disturbances are thought to exist across a wide range of chronic physiological and psychological conditions. Here we present a task and analysis procedure to quantify specific dimensions of breathing-related interoception, including interoceptive sensitivity (accuracy), decision bias, metacognitive bias, and metacognitive performance. Two major developments address some of the challenges presented by low trial numbers in interoceptive experiments: (i) a novel adaptive algorithm to maintain task performance at 70-75% accuracy; (ii) an extended hierarchical metacognitive model to estimate regression parameters linking metacognitive performance to relevant (e.g. clinical) variables. We demonstrate the utility of the task and analysis developments, using both simulated data and three empirical datasets. This methodology represents an important step towards accurately quantifying interoceptive dimensions from a simple experimental procedure that is compatible with clinical settings.

## Introduction

Understanding how the brain integrates sensory information to guide perception and action is a core component of neuroscientific research. Whilst the mapping of sensory pathways and perceptual phenomena have seen major developments in our understanding of the ‘exteroceptive’ domain (such as vision, audition, touch etc.), the study of ‘interoception’ (or the brain’s processing of sensory inputs from within the body) has begun receiving attention only relatively recently^1^. While theoretical concepts of the dynamic interplay of brain and body – including interoception, homeostatic and allostatic control^2–5^ – exist, empirical investigations have lagged behind. However, empirical studies of interoception have been recently boosted by a surge of interest in multiple neuroscientific fields, given that impairments in interoceptive processing have been proposed to play a role in emotions, decision making, consciousness and mental health^1,6^.

Perceptual processing is a complex phenomenon, and one that is highly integrated with other domains of brain function. For example, visual perception can be manipulated via changes in factors such as attention^7^, emotional state^8^ or expectation^9^. Furthermore, objective performance in perceptual detection tasks (i.e. accuracy and sensitivity towards stimulus detection, as often measured with classic psychophysics experiments^10^ and using inspiratory loading paradigms^11–15^) can be differentiated from more ‘metacognitive’ dimensions, where metacognition refers to the ability to accurately reflect and monitor cognitive or perceptual processes^15–19^. To quantify aspects of metacognition, measures of task performance can be paired with judgements of the confidence assigned to a decision^15,17,19,20^. From these metrics, average confidence can be thought of as a ‘metacognitive bias’, or a tendency towards a certain level of confidence, while ‘metacognitive performance’ (or ‘metacognitive sensitivity’) reflects how well confidence measures align with actual task performance^16–19^. Critically, to distinguish these metacognitive measures from underlying task performance, either objective accuracy needs to be held consistent across participants, or the effect of task accuracy needs to be accounted for using appropriate mathematical models (and an adequate volume of data acquired to fit these models)^17^. Here, we propose a method that incorporates both control over task performance, as well as accounting for any residual task performance variation between individuals.

These dimensions – task performance, metacognitive bias and metacognitive performance – have been distinguished within an interoceptive model by Garfinkel and colleagues^19^. Here, the authors demonstrated that these domains appear to be both quantifiable and distinct, and potentially related to traits such as anxiety. While their study was focused on cardiac-related body signals, initial work has hinted at potential cross-talk across different interoceptive ‘channels’ in the metacognitive domain, where corresponding interoceptive metacognition (but not task performance) was observed across cardiac and respiratory tasks^15^. Interestingly, this study also reported significantly elevated confidence in breathing-related perceptual decisions when compared to judgements of cardiac and tactile performance^15^. Whilst breathing is more consciously accessible for both perception and control than the cardiac domain, this elevated confidence also highlights the importance and relevance of breathing-related symptoms in the maintenance of homeostasis, whereby even a single breath of restricted or occluded breathing can be perceived as extremely unpleasant and frightening^21^.

Central for the further development of interoceptive research is the requirement to develop robust methodologies that can quantify interoceptive and metacognitive dimensions. Breathing is often considered to lie at the border of interoception and exteroception, combining cues from sensory avenues such as tactile and skeleto-muscular sensations across the chest wall, muscular effort, blood-gas signals representing bodily respiratory status, and air temperature and humidity, to name a few. Importantly, the accessibility of breathing to voluntary alterations and conscious perceptions lends itself to an array of experimental paradigms, including those that do not require exteroceptive cues. In a similar manner to cardiac measures, breathing contains inherent variability in flow and resistance both between and within individuals. These individual and breath-by-breath differences render highly accurate measures of breathing-related perceptual sensitivity challenging. However, if the performance of a perception task is both controlled and accounted for, metacognitive measures relating to interoception can become both accessible and independent of these challenges.

In this paper we provide a novel methodology for controlling task performance on a breathing perception task, and demonstrate the utility of applying a computational modelling approach to analyze metacognitive metrics of breathing perception. The main benefit of utilizing this computational model to assess metacognitive performance is that the effect of underlying task performance on metacognition can be removed^16,18^, as differences in task performance will produce concurrent differences in apparent metacognition if not adequately accounted for^17^. Importantly, an experimental setup is employed that is sufficiently simple and mobile to enable practical applications outside a laboratory setting, providing progress towards more useful clinical assessments of interoceptive properties of breathing. Inspiratory resistance is used as the breathing stimulus in this task, as it is both controllable and relevant to many individuals; for example, changes in airway resistance and pressure can result from both bronchoconstriction in conditions such as asthma and/or panic disorder^22^. However, inspiratory resistance also changes in physiological conditions, for example, simply as a result of increased inspiratory flow during activities such as exercise^23^ or hyperventilation induced by states of arousal^24^, but also as the result of reflex-mediated bronchoconstriction in response to cooling of the skin or upper airways^25,26^. Furthermore, it is now widely acknowledged that the perceptual system can be influenced by top-down factors such as attention, expectation and affect^3,27–36^; a known issue in conditions where symptoms are discordant with objectively measured medical markers, such as in asthma^34,37–39^ or those with medically unexplained symptoms^40–42^. Therefore, using a task that is able to dissociate measures such as perceptual sensitivity from decision bias and metacognition has great potential to fill an important unmet need in clinical practice.

To firstly demonstrate the utility of computational modelling for breathing, we combined an interoceptive breathing task based on resistive loads^15^ with an established computational model of metacognition (HMeta-d)^16–18^. We utilized both simulations and empirical data from individuals with asthma, as a cohort of individuals who regularly experience changes in airway resistance. The HMeta-d model utilizes a robust hierarchical statistical framework that allows computational models of metacognition, such as the meta-d model^16^, to be fit to task data where only low numbers of trials are available, such as those from interoceptive tasks. Furthermore, we present an extension to a hierarchical Bayesian model of metacognitive efficiency, HMeta-d^18^, that allows measures of metacognitive performance to be directly regressed against external variables of interest within the hierarchical model. This is an advantage over standard approaches as it capitalizes on the power of hierarchical estimation, especially when trial numbers are low, but avoids the problems encountered by post-hoc regressions on hierarchical model parameters such as unwanted shrinkage to the group mean. This shrinkage within standard hierarchical models is where individual subject estimates are drawn towards the group mean, and the variance between subjects (and thus the ability for post-hoc regression models to accurately explain inter-individual variance) is reduced. Lastly, we present an adaptive task-performance algorithm that directly targets a perceptual threshold accuracy of ∼70% and allows online control of performance, to aid the collection of a maximal number of trials at this level of task difficulty. Importantly, quantification of higher-order metacognitive measures is most efficient when objective performance is both significantly above chance (or guessing: 50%) and lower than a ceiling value of 100%, such that the resulting perceptual errors can be used to quantify the correspondence between objective accuracy and subjective confidence reports^17^. The utility of the analysis models and task-performance algorithm are established using both simulations and empirical data.

## Methods

The Methods section firstly contains an overview of how the Filter Detection Task (FDT) is carried out, followed by an explanation of the four interoceptive measures that can be quantified. We then describe the computational model simulations employed to determine the applicability of these analysis methods to limited-trial interoceptive applications. Next, we describe the testing and analysis methods employed to assess illustrative hypotheses in an example empirical dataset that encompassed a group of individuals with asthma as well as healthy controls. Finally, we describe the novel task algorithm that was designed to both control performance (within sessions and between individuals) and increase the number of trials measured at the perceptual threshold of an individual, which is defined using the classical value of 70-75%^43^, lying half-way between guessing between the two answer options available (50%) and a ceiling value of 100%. We use both simulations and two further empirical datasets to demonstrate the utility of the algorithm to control performance and reduce the number of excess (unused) trials.

### Filter Detection Task overview

To systematically test breathing perception within interoception, we have developed a perceptual threshold breathing task (the FDT) based on a previously-reported perceptual breathing task^15^. This task is a perceptual discrimination task, and can either be completed as a ‘Yes/No’ decision task, or a two-interval forced choice (2IFC) task. This task requires a computer to run MATLAB, as well as low-cost and easily accessible spirometry filters and anesthetic tubing, presenting a low-cost and simple setup (Figure 1) that allows for assessment of multiple interoceptive and metacognitive measures within the breathing domain. Participants wear a nose clip throughout the task, such that breathing is performed only through the mouth.

**Figure 1.**
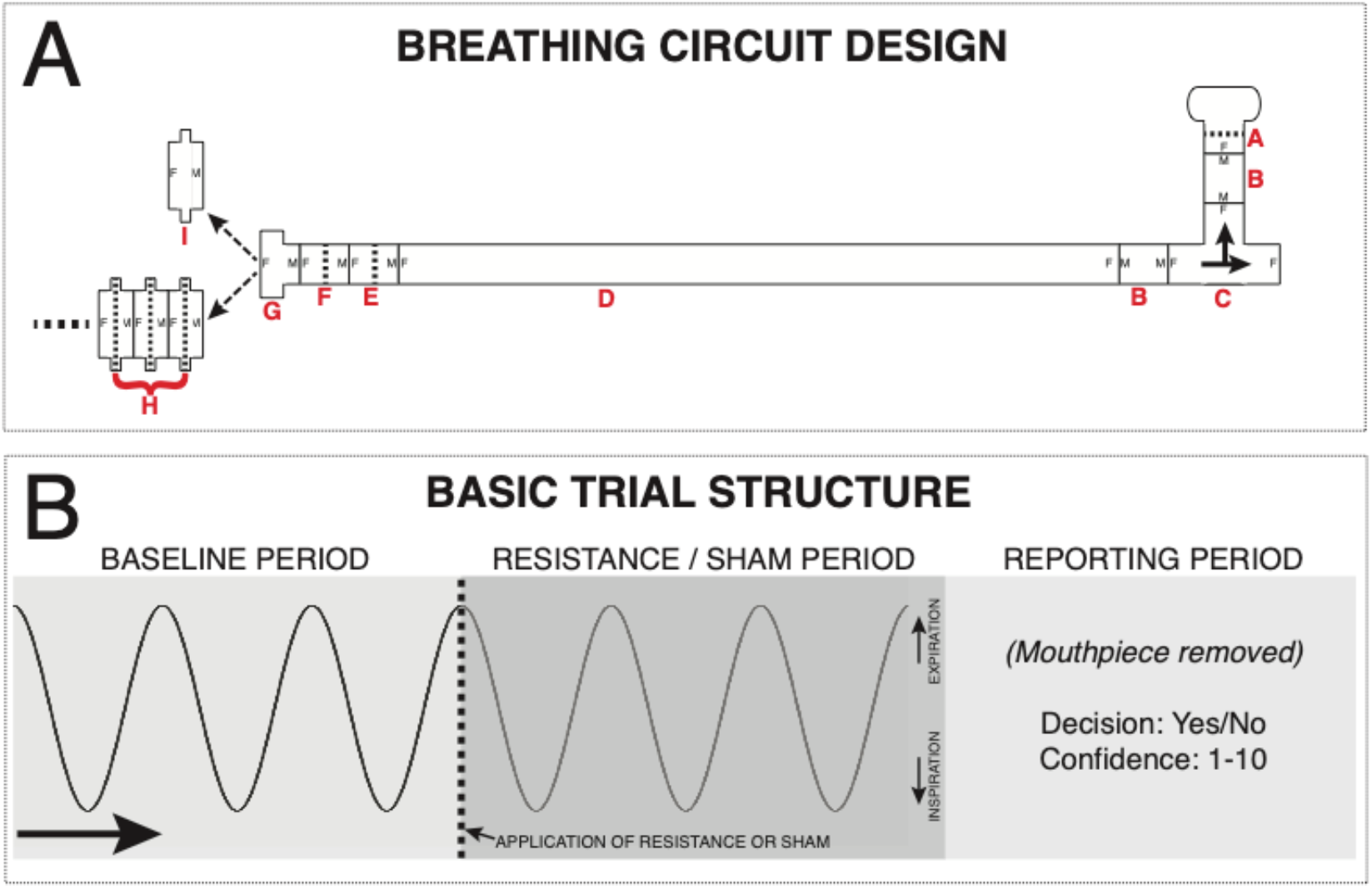
A) Diagram of circuitry for the filter detection task. A single-use, bacterial and viral mouthpiece (A: Powerbreathe International Ltd., Warwickshire, UK - Product SKU PBF03) is attached to a 22 mm diameter connector (B: Intersurgical Ltd., Berkshire, UK - Product 1960000) and a t-shaped inspiratory valve (C: Hans Rudolf, Kansas City, MO, USA - Product 1410/112622), connected to a 2 m length of 22 mm diameter flexible tubing (D: Intersurgical Ltd. - Product 1573000) and two additional baseline filters (E: Intersurgical Ltd. - Product 1541000, and F: GVS, Lancashire, UK - Product 4222/03BAUA). A 22-30 mm (G: Intersugical Ltd. - Product 197100) adapter then allows the attachment of either a series of connected spirometry filters (H: GVS - Product 2800/17BAUF, Pressure at 30 L/min < 0.3 cm H_2_O, Resistance < 0.48 cm H_2_O / L.sec^-1^) or a sham ‘dummy’ filter – a spirometry filter shell with the inner bacterial protection pad removed (I). B) Overview of the basic trial structure for a Yes/No formulation of the task. Participants take three normal size/pace breaths (with the sham filter attached), and during the third exhalation (indicated by the participant raising their hand and the dotted line in panel B) the experimenter either swaps the sham for a number of stacked filters (to provide a very small inspiratory resistance) or removes and replaces the sham filter. Following three more breaths, the participant removes the mouthpiece and reports whether they thought it a resistance was added (‘Yes’) or not (‘No’), and how confident they are in their decision on any scale (here 1-10 used, with 1 = guessing and 10 = maximally confident in their decision). If a two-interval forced choice (2IFC) formulation of the task is used, the filters (resistance) are either placed on the circuit for the first three breaths or the second three breaths according to the FDT algorithm, with the sham filter on the system during the alternate period. The reported decision from the participant is whether they thought the resistance was on in either the first set or the second set of three breaths, and also again the confidence in their decision.

For the ‘Yes/No’ version of the task, a standard trial structure consists of participants first taking three ‘baseline’ breaths through a mouthpiece connected to a simple breathing system (outlined in Figure 1). Following the baseline breaths, three breaths take place under ‘resistance’ or ‘sham’ conditions: either an inspiratory load is created via the addition of combinations of clinical breathing filters (signal trials, filters provided by GVS Filter Technology, product number 2800/22BAUF), or an empty filter (sham trials) is added to the system. The filters provide a resistance of < 0.48 cm H_2_O / L.sec^-1^ with the filter membrane attached (see Supplementary Material for further information). After each trial, participants are asked to verbalize their decision as to whether or not a load had been added. Alternatively, for the 2IFC task, the inspiratory load is either added in the first or second set of three breaths, and participants are asked to choose in which of the two intervals the inspiratory load was present. In either task, after each trial participants are asked to verbally report their confidence in this decision on a user-defined scale, e.g. from 1-10 (1 = not at all confident in decision, 10 = extremely confident in decision). The use of verbal feedback has the additional advantage of momentarily taking the participant’s attention away from their breathing. Participants can also take any length of rest period required between each trial.

The responses from the FDT can then be used to determine a variety of interoceptive measures (outlined below), including inspiratory load-related perceptual sensitivity, bias in symptom reporting, perceptual confidence and metacognition (the ability to accurately reflect upon cognitive or perceptual processes).

### Breathing-related interoceptive measures

In the original version of the filter detection task^15^, three domains of breathing-related interoception were quantified: interoceptive sensitivity (number of filters required for a discrimination threshold of approximately 75%), interoceptive sensibility (average confidence over threshold trials) and metacognition (correspondence between accuracy and confidence using a Type 2 receiver operating characteristic curve; an ROC curve). Here, we aimed to extend these measures to incorporate a more thorough and nuanced overview of a range of interoceptive dimensions. These measures include interoceptive sensitivity, decision bias, metacognitive bias, and metacognitive performance.

### Interoceptive sensitivity

This measure is analogous to interoceptive accuracy, and in this task aims to quantify the ‘perceptual threshold’ as a means for determining breathing-related interoceptive sensitivity. In other words, interoceptive sensitivity here is the level (i.e. number of filters) at which a participant is able to consciously detect an inspiratory loading stimulus. However, to allow for the quantification of higher-order interoceptive measures (such as metacognition), the number of filters used must elicit a performance that is both significantly above chance (or guessing) and lower than 100%, so that perceptual errors can be used to quantify the correspondence between accuracy and confidence (as explained below in ‘metacognitive performance’). To achieve a task difficulty that elicits this required performance, the original publication of this task^15^ utilized a descending accuracy staircase protocol, whereby a large starting filter number (for example 7 filters), and 20 trials were completed at descending filters until a final filter level when performance first dropped below 70%. While many staircase protocol options could be adopted to achieve a desired task difficulty, the time to complete one trial (approximately 30-60 seconds), the natural variability in resting tidal breaths^44^, the inherent bias associated with the Yes/No task formulation^45^ and the fixed filter intervals render many traditionally-employed staircase protocols (such as the two/three-down-one-up^10^) less straightforward with the current methodology. Therefore, we have developed a custom staircase algorithm (explained in ‘Task performance algorithm’) that employs probability metrics to objectively assess both task performance and trajectory towards the required perceptual threshold (see ‘Task performance algorithm’ section for full explanation of the algorithm).

### Interoceptive decision bias

If using the FDT as a Yes/No task, a quantifiable measure of behavior is the ‘bias’ towards reporting ‘Yes’ or ‘No’. This bias represents the placement of a criterion value above which the presence of a resistance is reported, reflecting an individual’s inherent tendency to report the presence of an inspiratory resistance. Importantly, this bias can be quantified using Signal Detection Theory (SDT^46,47^), and may represent an important cognitive trait regarding the experience of respiratory symptoms. Using SDT, stimulus sensitivity (d’) can be separated from bias (or the placement of a criterion, c)^46,47^. Using this theory, we are able to disentangle the components of measures that may be confounded by a mix of task sensitivity and bias, such as performance accuracy. When using the FDT as a 2IFC task, the measured ‘bias’ will instead be the tendency to report the resistance on the first or second interval. While this may have limited relevance to real world scenarios, quantification of d’ using this task design is likely to also allow for a more accurate representation of task sensitivity and a possibly more translatable measure of metacognitive performance^48^, which can both be confounded by variations in criterion placement.

### Interoceptive metacognitive bias

Average subjective confidence or metacognitive bias in interoceptive decisions has also previously been referred to as interoceptive “sensibility”^19^. In this task, we take an overall average of the confidence scores (measured across the perceptual threshold trials) to represent interoceptive sensibility that directly corresponds to the task at hand. Additional trait-like, global perceptual measures of sensibility could also be gathered by using separate interoceptive questionnaires such as the Porges Body Questionnaire^49^. Interoceptive sensibility is also referred to as “metacognitive bias”^17,18,50^, as it represents the tendency to give higher or lower confidence ratings.

### Interoceptive metacognitive performance

Breathing-related metacognitive performance (also termed ‘interoceptive awareness’^19^ and ‘interoceptive insight’^1^ in the literature) in this instance is considered to be the correspondence between task accuracy and confidence^19^, or the ability to recognize successful perceptual processing^17^. Previous reports of interoceptive metacognition have utilized the area under a type 2 ROC curve^15,19,20^, resulting in a measure of *absolute* metacognition where the effect of underlying task performance also influences the final score^17^. However, more recent model-based approaches have developed a metric of metacognitive performance that can subsequently take into account task performance, known as meta-d’^16^. Meta-d’ represents “the sensory evidence available for metacognition in signal-to-noise ratio units”^17^, which, because it is in the same units as d’, can be straightforwardly compared to task performance as a ratio (Mratio: meta-d’/d’; the log of the ratio, logMratio, is also often used to meet Gaussian assumptions: log(meta-d’/d’)). This ratio of absolute metacognitive performance (meta-d’) divided by task performance (d’) is thus a *relative* measure of metacognitive performance (often termed ‘metacognitive efficiency’). In order to employ this model-based approach for the FDT, a hierarchical formulation of the meta-d model (HMeta-d) is employed that allows efficient pooling of data from multiple subjects^18^, allowing us to estimate model parameters on a relatively small number of threshold trials (>= 40 trials).

### Metacognitive model simulations

To demonstrate the feasibility of utilizing the meta-d model for interoceptive tasks with low trial numbers, we first present simulated results to establish the recoverability of group metacognitive performance (Mratio) values using the original maximum likelihood estimation algorithm (MLE^16^), a single-subject Bayesian model and a hierarchical (group) Bayesian model (HMeta-d), which are both provided in the HMeta-d toolbox^18^. Parameter inference in the HMeta-d toolbox rests on a Markov chain Monte Carlo (MCMC) sampling procedure, implemented using the JAGS software package (http://mcmc-jags.sourceforge.net). The extent of the group Mratio recoverability is demonstrated for 20, 40 and 60 trials per subject, with 30 simulated participants. Simulations were generated from a set of values *N*(*μ, σ*), which refer to Gaussians parameterized by a mean and standard deviation, using the metad_sim function provided in the HMeta-d toolbox^18^. The meta-d’ values for the first set of simulations were generated from seven group Mratio distributions (meta-d’ / d’) with parameters *N*([0.25 0.5 0.75 1.0 1.2 1.5 1.75 2], 0.1), where d’ ∼ *N*(1, 0.1) and c ∼ *N*(0, 0.1).

Second, we developed and simulated an extension of the HMeta-d model (RHMeta-d), which incorporates a simultaneous hierarchical estimation of a regression parameter (beta) that controls variation in logMratio values in relation to a subject-level predictor (such as a clinical score). The model was adjusted as follows (for full details of the original model please see the original publication^18^). *N*(*μ, σ*) and *HN*(*μ, σ*) refer to Gaussians and Half-Gaussians parameterised by a mean and standard deviation, while *T*(*μ, σ, v*) refers to a T-distribution parameterised by a mean, standard deviation and degrees of freedom:

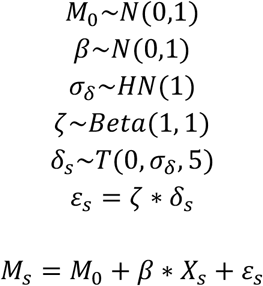

where *M*_*s*_ refers to the log(meta-d’/d’) value for subject *s, M*_0_ is the baseline logMratio for the group (i.e. the intercept of the regression), and *X*_*s*_ is a vector of predictor values (e.g. clinical scores) for each subject. This formulation embeds the estimation of psychopathology-cognition relationships into the parameter inference routine, such that the group-level posterior over *β* reflects the influence of individual differences in *X* on metacognitive performance^51^. To ensure that the regression is robust to outliers, the noise *ε*_*s*_ is drawn from a T-distribution with a standard deviation of |*ζ*|*σ*_*δ*_ and 5 degrees of freedom^52^. Consistent with the original HMeta-d model^18^, a redundant multiplicative parameter *ζ* is used to introduce an additional random component in the sampling process to aid the recovery of the posterior on the noise scale.

For these simulations, group baseline logMratio (*M*_0_) values were generated from a distribution ∼ *N*(log(0.8), 0.1). Then, to simulate data from each subject, a random value of the covariate *X*_*s*_ was drawn from a standardised distribution *N*(0,1), multiplied by a group regression coefficient that was one of a fixed set of β values (β ∼ [-0.5:0.5]) and added to this baseline logMratio together with zero-mean noise sampled from *N*(0, 0.1). Values used for d’ and c’ were consistent with previous simulations (d’ ∼ *N*(1, 0.1) and c ∼ *N*(0, 0.1)), and data were generated using either 20, 40 or 60 trials per subject and a confidence scale of 10 rating points. From these simulated data, parameter estimates were then obtained either using the original HMeta-d model combined with a post-fit linear regression (i.e. a standard linear regression conducted on the per-subject point estimates obtained from the group-level fit, which we denote HMeta-d+R), as well as the extended hierarchical regression model (RHMeta-d) described above. An average for ten sets of simulations for each group β value was calculated for the final results.

### Analysis methods: Example dataset of asthma and healthy controls

To demonstrate the utility of the analysis methods, we employed an example dataset that included a group of individuals who experience an elevated frequency of breathing symptoms. While it has been observed that individuals with asthma can vary from under-reporting to over-reporting of symptoms^34,37–39^, group-wise analyses of asthma have demonstrated both an elevated prevalence of anxiety and depression symptoms^53^ and that symptom prevalence is related to these affective qualities^33,54–56^. Furthermore, as the metacognitive properties of individuals with asthma have not yet been systematically tested and could viably relate to symptom reporting, this group was selected as an example test-case for this method. Importantly, the FDT allows us to separate the effect of interoceptive sensitivity to inspiratory loads, bias towards under- or over-reporting the presence of a resistance, metacognitive bias (higher or lower confidence in interoceptive decisions) and metacognitive performance (‘insight’ into perceptual performance). These separable entities may help to shed light on the potential drivers towards the under- to over-reporting of symptoms^34,37–39^ in asthma.

Sixty-three individuals with asthma (39 females, mean age (± sd) 43.7 ± 12.2 years, recruited through general practitioner clinics and public advertisements) and 30 healthy controls (19 females, mean age (± sd) 44.2 ± 12.2 years, recruited through public advertisements) took part in a study approved by the Oxfordshire Clinical Research Ethics Committee. Participants underwent the FDT and completed the Dyspnea 12^57^ questionnaire as a subjective assessment of their breathlessness severity. Additionally, participants completed a further set of questionnaires and additional physiological and behavioural measures that will be addressed elsewhere. Seven individuals with asthma were excluded from the analysis due to insufficient data (10 trials or less of the FDT, n = 4), or performance of less than 50% correct (n = 3), as determined in the preregistered analysis plan (https://gitlab.ethz.ch/tnu/analysis-plans/harrisonetal_fdt_methods_2020).

During the FDT in this study, the number of filters required for each participant to induce task performance at perceptual threshold was determined manually, using a staircase method adjusted from a previous publication of the task^15^ (data was collected prior to the development of the task algorithm). In this step-wise procedure, participants first completed 10 trials at 4 filters. If the task accuracy was below or above 70%, the number of filters was adjusted up or down accordingly by one filter. Performance accuracy was assessed again at 20, 30 and 40 trials, with adjustments made if the accuracy moved outside of the 65-75% range (with an acceptable range of 60-80% in later trials). The aim was to complete 40-60 trials, which was limited by time and attention constraints of each participant. A 0-100 confidence rating scale was employed, with 0 = not at all confident, and 100 = maximal confidence. These confidence scores were down-sampled into 10 rating bins prior to analysis with the HMeta-d model.

To demonstrate empirical questions that could be answered using the FDT and metacognitive models, we firstly tested any differences between individuals with asthma and healthy controls across all FDT parameters (https://gitlab.ethz.ch/tnu/analysis-plans/harrisonetal_fdt_methods_2020). For the measures of interoceptive sensitivity (number of filters), decision bias (*c* parameter from model) and metacognitive bias (average confidence), tests for data normality were first conducted using Anderson-Darling tests, with an alpha value of p < 0.05 required to reject the null hypothesis of normally distributed data. A significant group difference was then tested using two-tailed parametric or non-parametric Wilcoxon rank-sum tests. A group difference in metacognitive performance (the Mratio parameter from the HMeta-d model output) was assessed by first calculating the distribution of differences in posterior parameter samples from each group (control Mratio samples > asthma Mratio samples), and then determining the highest density interval (HDI) for this distribution. The HDI employed was a two-tailed 99% (Bonferroni corrected for five tests) confidence interval, where a significant difference between groups was denoted if the resulting HDI did not span zero. Significance for all other tests was denoted by p < 0.01 (p < 0.05 Bonferroni corrected for the five experimental tests).

To assess a possible relationship between metacognitive performance and the prevalence of reported breathing-related symptoms, a direct test of the relationship between metacognitive performance and D12 was performed using a linear regression model in the individuals with asthma. To achieve this, the HMeta-d model was extended to include a hierarchical estimation of a linear regression parameter, whereby a group regression coefficient (beta) was simultaneously fit within the model to determine the relationship between logMratio and D12. The significance of the beta parameter was then also determined using its posterior samples, with a two-tailed 99% HDI (Bonferroni corrected) that did not span zero determining a significant relationship between more severe D12 and worsened metacognitive performance. An illustrative procedure using a split-half analysis is presented in the Supplementary Material.

### Task performance algorithm: Simulations and empirical data

Lastly, to assist in the collection of a greater number of usable trials for further instances of the FDT, we created a novel staircase protocol (within a MATLAB toolbox package) to aid the selection of the appropriate number of filters for each participant. While adaptive psychophysics staircase algorithms are available (such as QUEST^43,58^), many of these formulations rely on adjustable and small available step-sizes and small amounts of sensory noise, and also often assume a pre-determined psychometric for model fitting. Therefore, for this novel application in the breathing domain we designed an algorithm that does not assume any psychometric function; instead, it estimates the underlying accuracy that gives rise to the current performance using a beta-binomial model. This simple model is robust and does not run the risk of non-convergence as can be observed with a more complicated algorithm such as QUEST, which may occur due to both the limited number of trials, step sizes available and variability in breathing from trial to trial.

When interfacing with the task algorithm, the researcher is given instructions for each trial via the MATLAB command window. Additionally, the researcher is required to enter the participant’s decision and confidence scores into the MATLAB command window when prompted for every trial, and this information is then used to dynamically update the staircase procedure. This staircase begins with one or two practice trials and a short calibration, which are completed under the same within-trial format (i.e. Yes/No or 2IFC structure) as the main task trials. In the practice trials for the Yes/No task, an additional ‘explicit dummy’ is first applied, whereby participants are told that it is a dummy. Another practice trial is then performed using a large load of 7 filters, where no feedback is given (no feedback is maintained for the rest of the protocol). The practice is then immediately followed by the calibration trials, where participants complete trials that increase by one filter each trial (beginning with a dummy) until they have correctly reported the resistance for two consecutively increasing filter numbers. A final calibration trial is then given, where the number of filters is dropped by one from the last trial. If participants correctly report the final calibration trial, they begin on that number of filters, whereas if they are incorrect, they begin with one additional filter. A diagram of the basic trial structure is presented in Figure 1, and the practice, calibration and real trial trajectory is provided in the Supplementary Material (Supplementary Figure 1).

Once the calibration is complete (or alternatively, a manual starting point can be provided), the main task trials begin. The target number of trials is specified (recommendation of >= 60 trials), and a pseudo-randomized sequence of trials are presented (trials are balanced between present/absent for a Yes/No task or between first and second interval for 2IFC). The target is for participants to be within a 65-80% accuracy band. Given a set of binary trials (and an appropriate prior), we use the fact that the posterior distribution over the underlying accuracy follows a beta distribution. After 5 trials at one filter level (with at least one resistance present for the Yes/No task), the posterior probability that the underlying performance accuracy for the current task difficulty (i.e. the current number of filters) falls between 65-80% is calculated using the difference in beta cumulative distribution functions for 80% and 65%, in a similar vein to the QUEST algorithm^43^. We use a weak prior on the accuracy itself (beta distribution with the parameters α = 2 and β = 1; prior mean = 67% accuracy, interquartile range = 37%), determined by the performance of simulations to produce observed accuracies closest to 75%. If the probability that the underlying accuracy is between 65-80% falls below a threshold of 20%, an addition or removal of a filter is automatically suggested to decrease or increase task difficulty respectively. If a new filter number is started, the trial count will begin again and 5 trials (with at least one resistance) must be completed before the algorithm will suggest any changes. If the filter change moves the filter number back to that of previous trials, the trial count will pick up again from the last trial at this level. In this instance, 3 trials (with at least one resistance) must be completed before any changes are suggested.

To demonstrate the utility of this task performance algorithm, we firstly present simulation results from a range of possible participant performances, characterized by a distribution of potential psychometric functions. These psychometric functions (n = 350) were constructed from an underlying logistic sigmoid with a lower asymptote at 0.5 (to account for chance answers with the two answer options of ‘yes’ and ‘no’), a slope *k* = [0.7:1.2], the number of filters at which the 75% threshold is obtained t = [1:7] and added Gaussian noise ε = *N*(0, [0.05:0.015]). We then ran each of the sigmoids generated from each of 5 starting points – from two filters below to two filters above the t parameter, totaling 7000 simulations. Second, we provide data metrics (number of trials, performance accuracy, number of filters) for two collected datasets using the Yes/No version of the task. The first of these collected datasets stems from the first 50 participants measured as part of a wider study approved by the Cantonal Ethics Committee Zurich (Ethics approval BASEC-No. 2017-02330). For this study, we employed a ‘constant’ staircase formulation of the algorithm, with the aim of collecting 60 trials at a single level of filters that elicited a task performance between ∼60-85%. The second empirical dataset includes the first 22 participants measured as part of a wider study approved by the Cambridge ethics committee (Ethics approval PRE.2018.092). This study employed a ‘roving’ staircase with 60 trials total, where the aim was to simply collect 60 trials regardless of the number of filters. In this scenario, the ‘threshold’ filter becomes the average of the number of filters employed across the task.

## Results

The Results section firstly outlines and compares the computational model simulations using three different implementations of the Metacognitive (Mratio) model. The hierarchical version (HMeta-d) is convincingly shown to be the most reliable in recovering simulated values of Mratio. Simulation results also establish the recoverability of regression parameters using the extended RHMeta-d model compared to the standard HMeta-d model. We then present the results from the example empirical analyses proposed in individuals with asthma and healthy controls, to demonstrate how the model outputs can be interpreted in light of example hypotheses. Lastly, we ascertain the utility of the novel task algorithm using both simulated and empirical results.

### Metacognitive model simulations

The simulation results firstly demonstrate that utilizing the hierarchical Bayesian HMeta-d model fit allows adequate recovery of group Mratio values (Figure 2). This recovery is possible even using as few as 20 trials per subject, with slightly larger uncertainties (demonstrated by the width of the highest density intervals) than those obtained for 40 and 60 trials. It is instructive to compare this to the alternative estimation methods: while MLE is able to recover an average Mratio value that is indeed representative of the simulated value, the uncertainty around these estimates (demonstrated by the width of the confidence intervals) is large when using even 60 trials per subject. Moreover, the confidence interval around these MLE estimates incorrectly encompasses zero for all group Mratio values below 1. The recovery of the group Mratio using the Bayesian single subject fit also has large uncertainties and shows shrinkage effects towards zero, recovering Mratio values below the simulated values across all trial numbers tested here.

**Figure 2.**
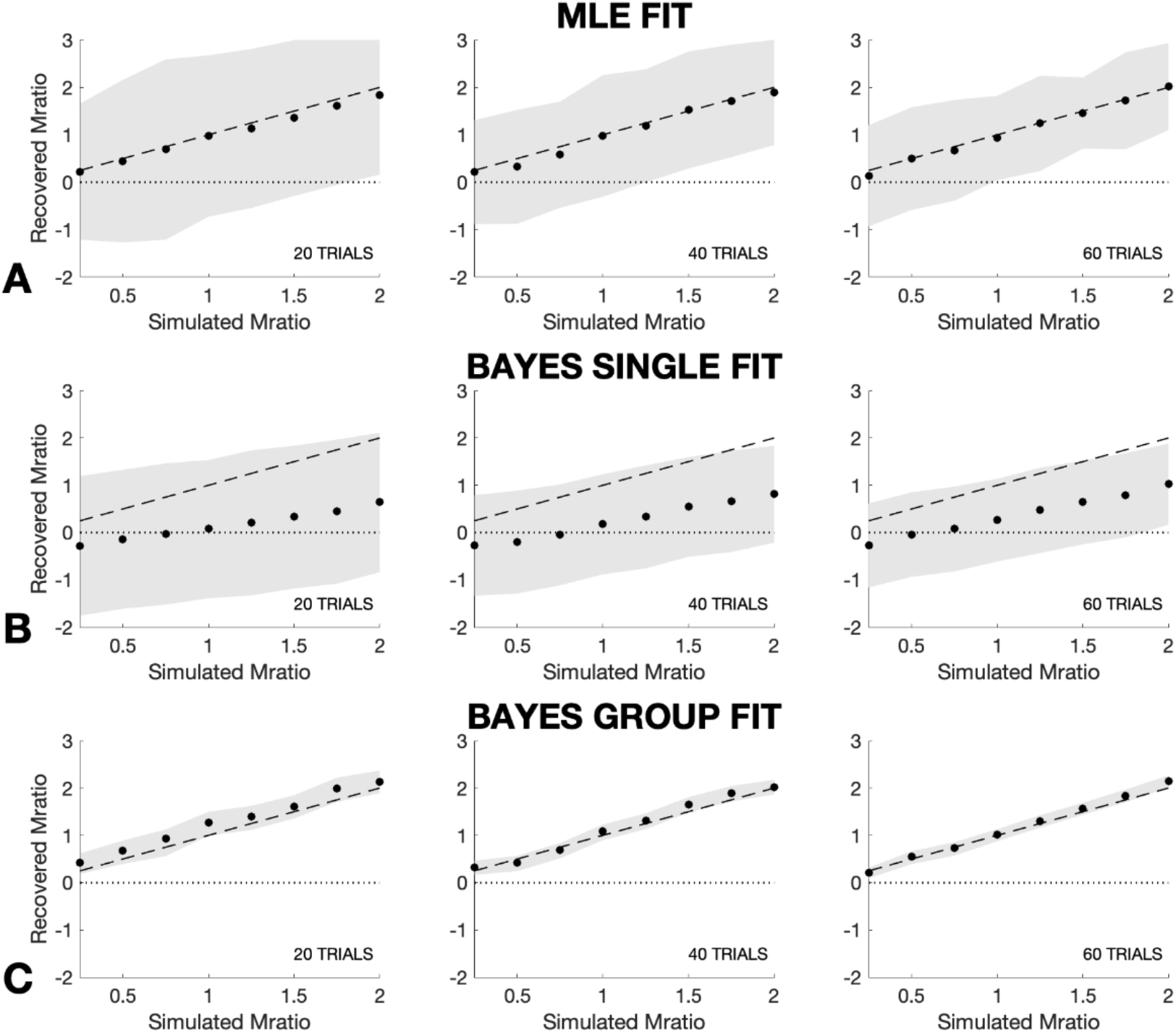
Group Mratio recovery for 20, 40 and 60 trials using three different meta-d models. Data were simulated from 8 groups of 60 subjects with group mean Mratio (meta-d’ / d’) values set to [0.25 0.5 0.75 1.0 1.25 1.5 1.75 2] ± 0.1 (sd). All simulated values were generated from data where d’ ∼ N(1, 0.1) and c ∼ N(0, 0.1), and a confidence scale of 10 rating points was used. A) Simulated vs. recovered Mratio values using maximum likelihood estimation^16^, where the shaded grey areas denote the 95% confidence interval of the estimate. B) Simulated vs. recovered Mratio values using a Bayesian single-subject fit (provided in the HMeta-d toolbox^18^), where the grey areas denote the 95% highest density interval (equivalent to a 95% credible interval) of the sampled estimate. C) Simulated vs. recovered Mratio values using a hierarchical Bayesian group fit (provided in the HMeta-d toolbox^18^), where the grey areas denote the 95% highest density interval of the sampled estimate. Dashed lines represent ideal recovery, with dotted lines at zero demonstrating the ability of the model fit to significantly recover group estimates (i.e. when confidence or highest density intervals do not include zero).

The second set of simulations were designed to probe the recoverability of single-subject Mratio values, for possible use in analyses comparing individual metacognitive performance against an external variable. Using the original HMeta-d model, we demonstrated that a post-hoc regression on the single-subject values was unable to accurately recover a group regression parameter simulated from the range β = [-0.5:0.5] even when using 60 trials, with all confidence intervals on the regression parameters encompassing zero (Figure 3). This is unsurprising given that the hierarchical model naturally shrinks single-subject estimates towards the group mean, losing information about individual differences. In contrast, the R-HMeta-d model was able to significantly recover beta values of ± >0.2 with 60 trials, ± >0.25 with 40 trials, and ± >0.3 with 20 trials (Figure 3). The results for multiple regression models (with up to three covariates) at each of 20, 40 and 60 trials are presented in the Supplementary Material (Supplementary Figure 5).

**Figure 3.**
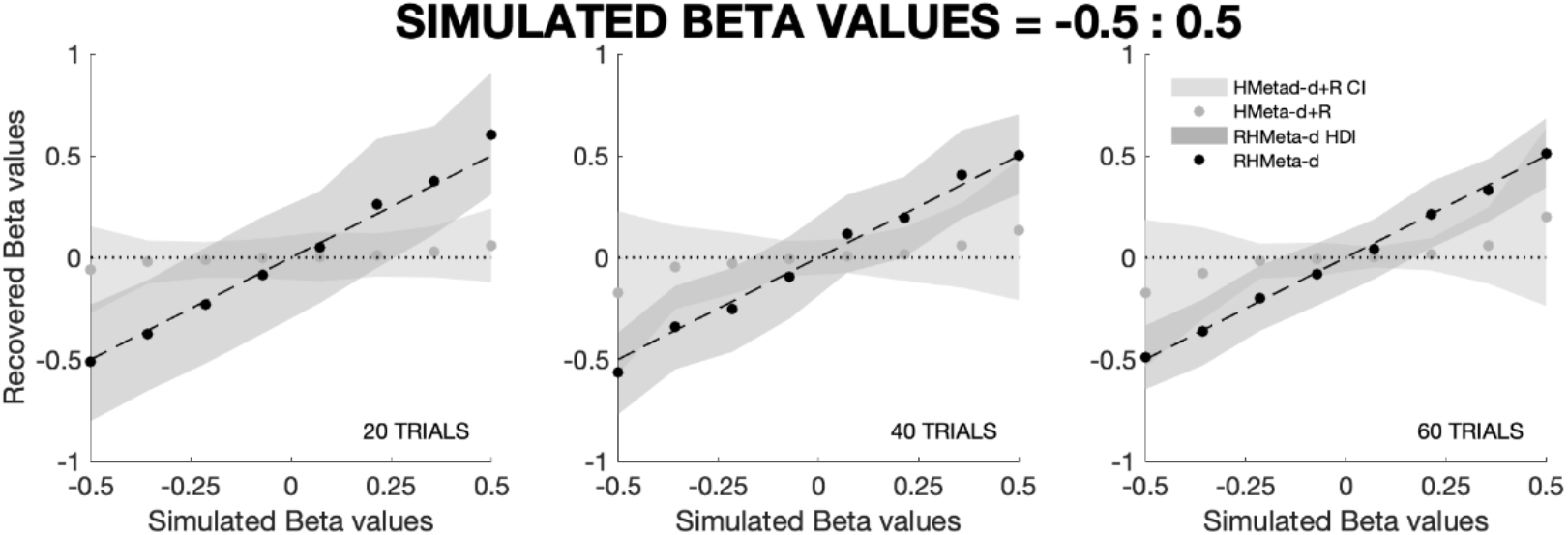
Demonstration of the recovery of group regression parameters (β ∼ [-0.5:0.5]) using either the original HMeta-d model combined with a post-fit linear regression (HMeta-d+R), or the extended regression HMeta-d (RHMeta-d) model. Ten sets of simulations were performed and results were averaged, where each simulation set included 60 simulated ‘subjects’ where logMratio = logMratio_baseline_ + β*covariate + noise, where logMratio_baseline_ ∼ N(log(0.8), 0.1), covariate ∼ N(0, 1), β ∼ [-0.5:0.5], noise ∼ N(0, 0.1), and with d’ ∼ N(1, 0.1), c ∼ N(0, 0.1). Grey areas denote the 95% highest density interval of the sampled estimate. Dashed lines represent ideal recovery of group beta values, and dotted lines at zero demonstrate the ability of the model fit to significantly recover group estimates (i.e. highest density intervals that do not including zero).

### Empirical data analyses

When considering the comparisons between the asthma and control groups, no significant difference was found between the two groups for interoceptive sensitivity (number of filters = mean ± se: controls = 2.87 ± 0.29, asthma = 2.80 ± 0.20), decision bias (c = mean ± se: controls = 0.01 ± 0.07, asthma = -0.01 ± 0.06), metacognitive bias (average confidence % = mean ± se: controls = 66.57 ± 2.99, asthma = 69.74 ± 2.13), nor metacognitive performance (Mratio = mean ± se: controls = 0.83 ± 0.14, asthma = 0.79 ± 0.12) (Figure 4). We then considered the relationship between breathing symptoms and metacognition in asthma only. While the data demonstrated a tendency for reduced metacognitive performance with higher symptom loads using a hierarchical regression analysis (RHMeta-d; Figure 5), which was not statistically significant (determined by an HDI that does not encompass zero). Using a hierarchical regression approach, the beta parameter mean was estimated as -0.22 ± 0.16 (se), with the beta HDI in the range [-0.67, 0.27] (Figure 5).

**Figure 4.**
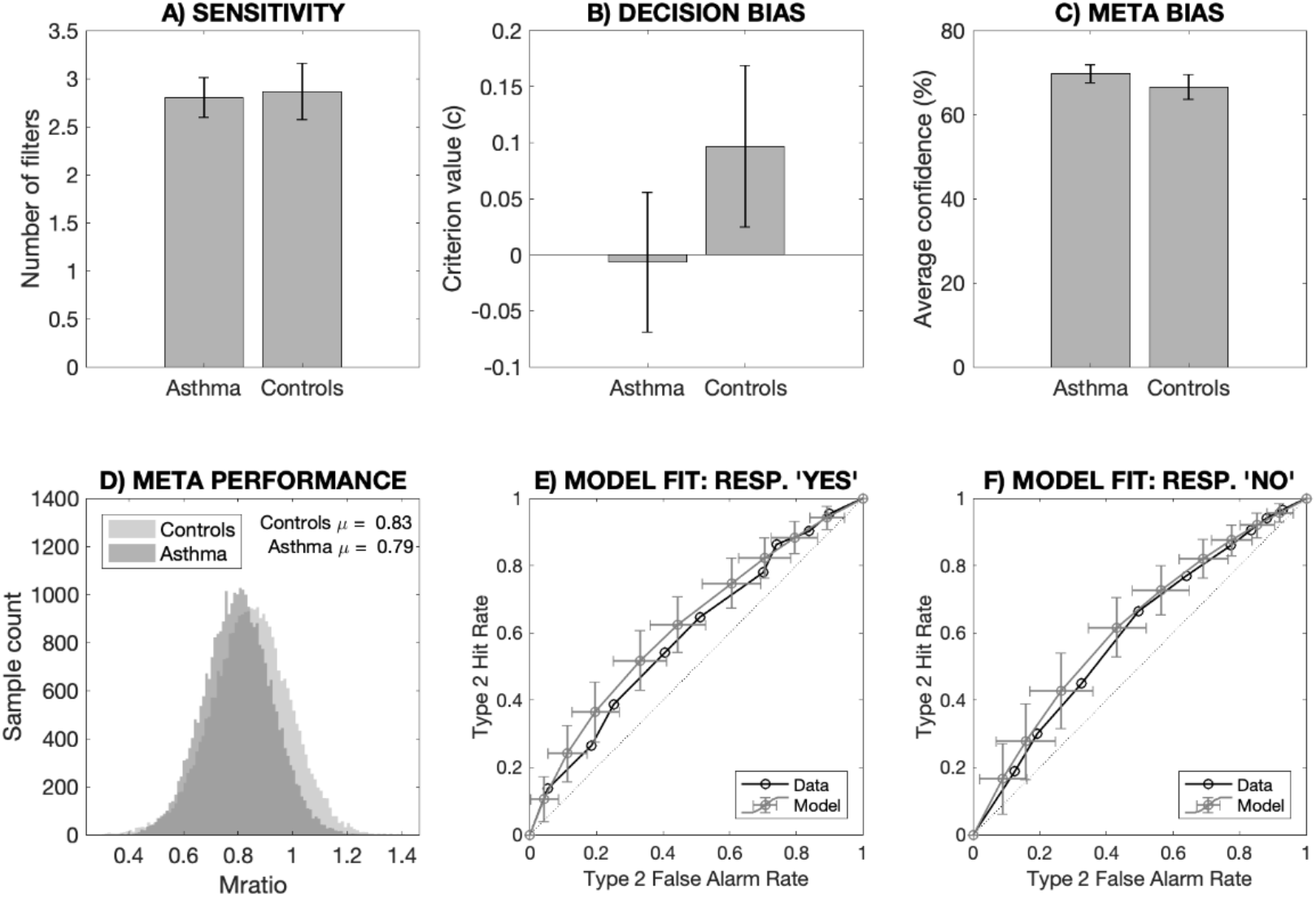
Group difference comparisons between individuals with asthma and healthy controls. Panels A, B and C denote group mean and standard error for each group regarding interoceptive sensitivity, decision bias and metacognitive bias, with no significant differences found between the groups. Panel D demonstrates the sampled posteriors for the group estimates of metacognitive performance (Mratio). Panels E and F demonstrate the model fit by comparing the observed and model estimates of the Type 2 ROC curves for both ‘Yes’ and ‘No’ responses (regarding the presence of an added inspiratory resistance). Model fits for all other models are presented in the Supplementary Material (Supplementary Figure 6).

**Figure 5.**
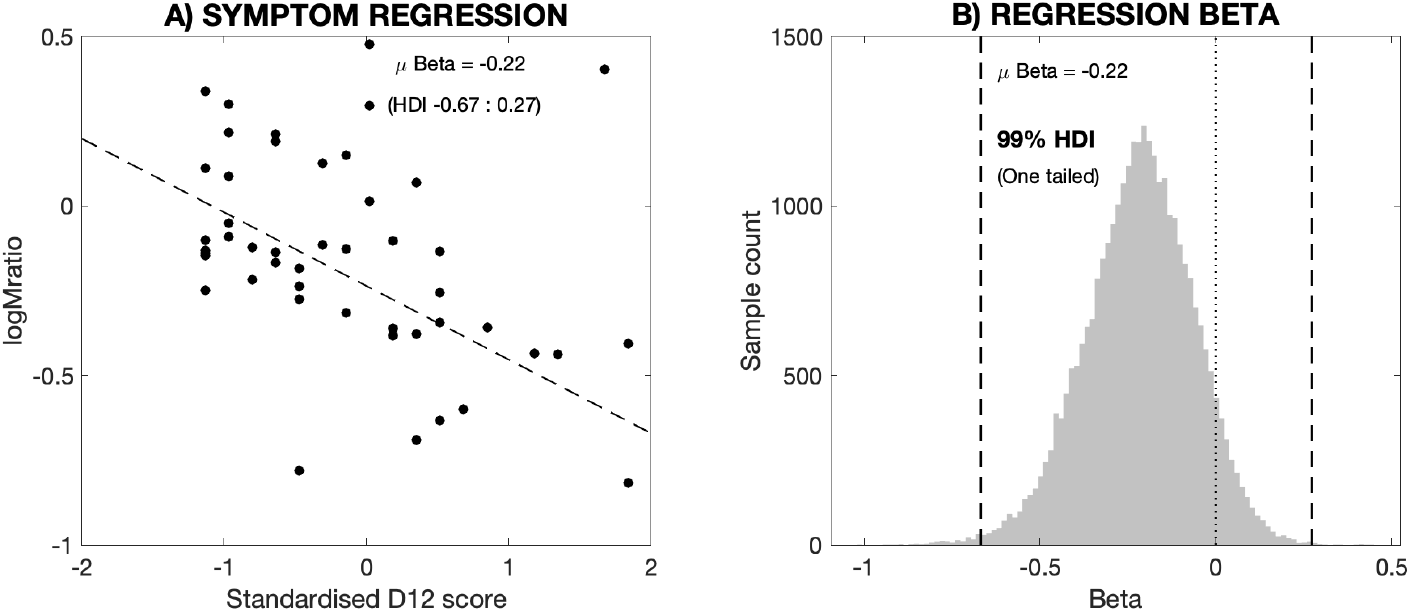
Comparisons between metacognitive performance (logMratio) and symptom load in asthma, using a hierarchical regression analysis. A) A hierarchical regression predicting logMratio from the standardised D12 scores within asthma participants. The regression was fit using an extension of the HMeta-d model (R-HMeta-d) in which the beta regression coefficient was fit simultaneously together with the logMratio scores. Dashed line represents the regression line from the model fit. B) The distribution of samples over the regression beta parameter (from panel A) fit using the R-HMeta-d model. Dashed lines represent the two-tailed 99% HDI which encompasses zero (dotted line) consistent with no significant relationship between D12 score and metacognitive efficiency in this dataset.

### Task performance algorithm: Simulations and empirical data

The results from both simulated and empirical data demonstrate the ability of the task algorithm to target performance towards a perceptual threshold that lies above chance (50%) and below ceiling (100%) performance (Figure 6). Simulations conducted using the task algorithm (with a ‘constant’ staircase formulation) produced task accuracy scores with a mean of 74.1 ± 8.7% (sd) and an accurate recovery of the 75% filter number, irrespective of the starting filter value (Figure 6A). Empirical data collected using the Yes/No formulation of the task (with a constant staircase) produced a task accuracy with a mean of 68.9 ± 7.4% (sd), with the threshold filter number spread between 1 and 8 filters (Figure 6B). Both the simulations and real data demonstrate a feasible number of trials (70.4 ± 10.3 (sd) trials for simulated results with 60 threshold trials, 69.6 ± 10.0 (sd) trials for empirical data) required to complete 60 trials at the threshold filter. In real terms, this indicates that it is possible to reliably measure respiratory interoception and metacognition in approximately one hour or less (assuming approximately 45-60 seconds to complete each trial). These estimates include a constraint whereby the algorithm was additionally programmed to continue until 30 trials are completed at the threshold filter number, followed by the option for manual intervention to instigate filter changes every ten trials if the accuracy moves out of acceptable bounds (experimenter decision required, depending on time taken and participant).

**Figure 6.**
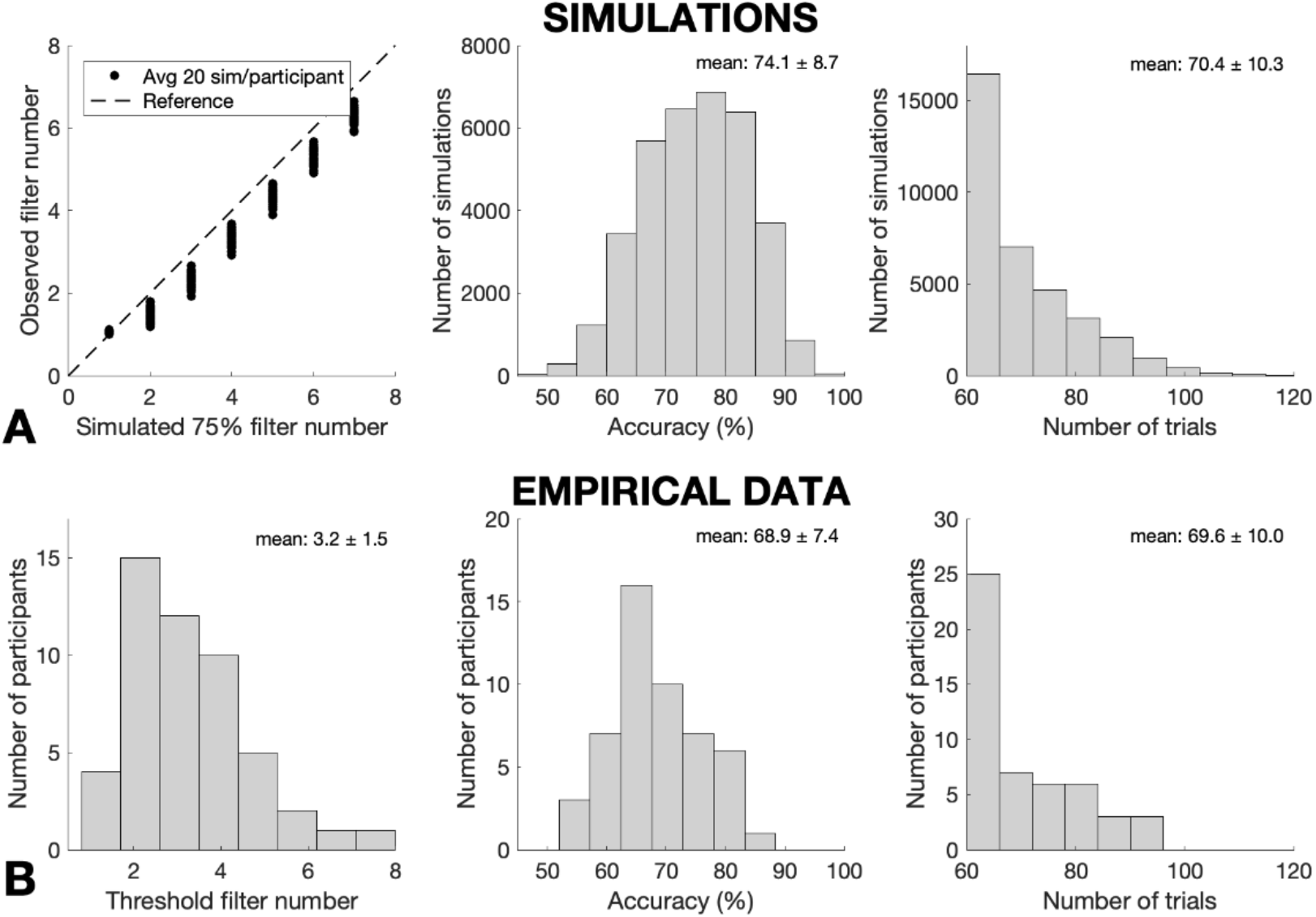
Results demonstrating the use of an adapted staircase algorithm for targeting a specified level of task difficulty over 60 trials. A) Simulation results, where data were generated from a range of logistic sigmoid functions bounded between 0.5 and 1, with 20 simulations for each sigmoid (‘participant’) from each of five starting points – from two filters below to two filters above the 75% threshold filter. Left: Simulated and recovered 75% filter number for each simulated ‘participant’. Middle: Histogram of the task accuracy scores for the 60 threshold trials for all simulations. Right: Histogram of the total number of trials required to complete 60 threshold trials for each simulation. B) Data collected using a Yes/No version of the task (with a constant staircase), where 50 participants each completed 60 threshold trials. Left: Histogram of the measured threshold filter number for each participant. Middle: Histogram of the task accuracy scores for the 60 threshold trials for the 50 measured participants. Right: Histogram of the total number of trials required to complete 60 threshold trials for each participant. All histograms are reported with mean ± standard deviation.

When utilizing a ‘roving’ staircase experimental design, simulations of the task algorithm produced task accuracy scores with a mean of 75.4 ± 8.0% (sd) and an accurate recovery of the 75% filter number, irrespective of the starting filter value (Supplementary Figure 8A). When the first 60 trials from the constant staircase empirical data discussed above were analyzed as a roving staircase (i.e. the first 60 trials analyzed, regardless of filter intensity), the mean task accuracy was slightly reduced from 68.9% to 67.7%, and the variance of the scores increased from 7.4% to 8.5% (sd) (Supplementary Figure 8B). A final dataset, collected with explicit use of the roving staircase paradigm, demonstrated a mean accuracy of 69.7 ± 11.7% (sd), and threshold filter numbers that ranged from 1-8 filters (Supplementary Figure 8C). No additional trials are required when utilizing a roving staircase design, as all trials following the calibration step are included in the analysis. The simulated and empirical results from the calibration algorithm are presented in Supplementary Figure 9.

Direct comparison between the empirical data collected using the three methods (i.e. manual accuracy calculations every 10 trials, the constant staircase design and the roving staircase design) is provided in Figure 7. Data collected using a manual accuracy calibration (asthma and controls) produced task accuracy scores with a mean of 66.4 ± 8.2% (sd), with a large number of additional trials (percentage of the number of threshold trials collected = 71.9 ± 17.7% (sd)) and two participants whose task accuracy was ≤ 50%. No difference in overall accuracy was found (Wilcoxon rank-sum tests) between any of the task designs (all p > 0.05), however the roving task design produced the largest standard deviation in the task accuracy across the methods (observed in Figure 7). Additionally, while a significant number of additional trials were required when using the manual and constant staircase methods (Wilcoxon signed-rank tests, p < 0.001 for both tests against zero), the constant staircase significantly reduced the additional number of trials required from the manual method, both as an absolute number of trials and as a percentage of the number of threshold trials collected (Wilcoxon rank-sum tests, p < 0.001 for both tests).

**Figure 7.**
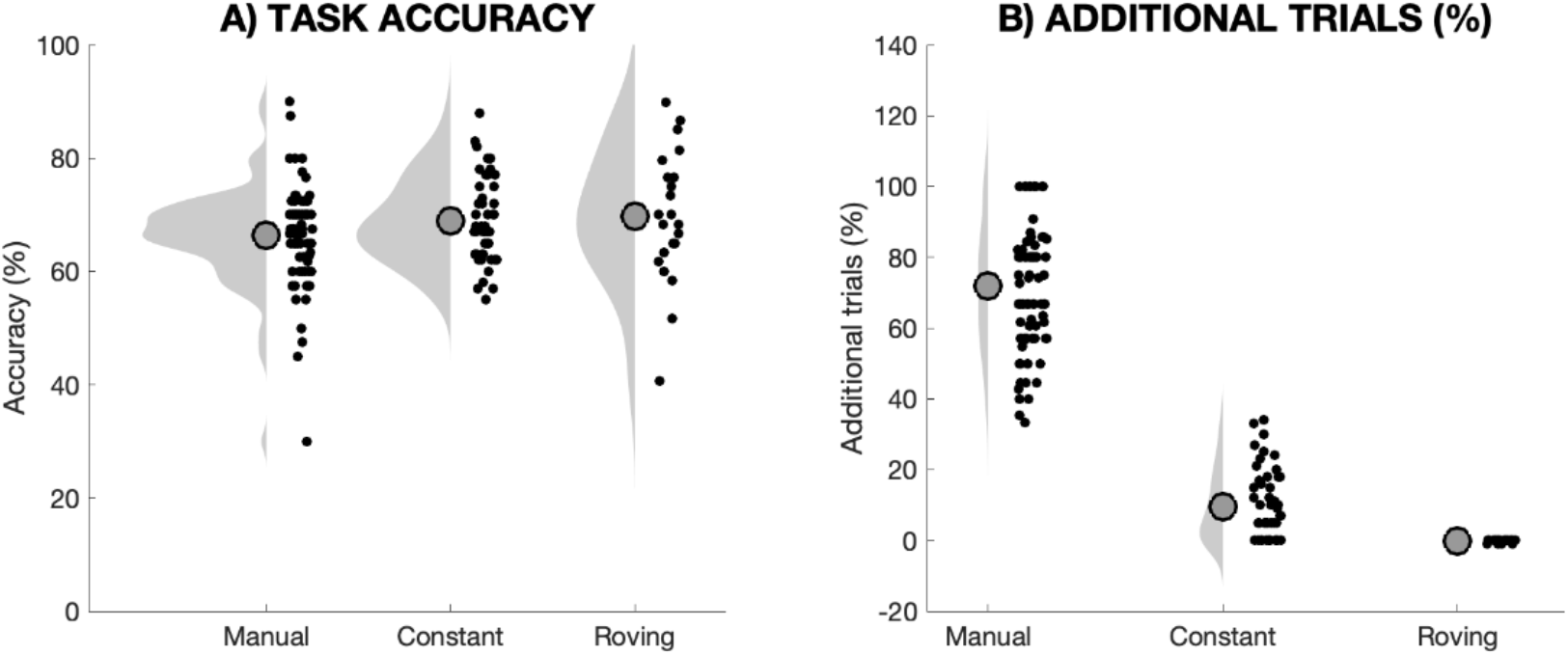
Comparison of the three empirical datasets collected using different methods: ‘Manual’ = Manual staircase adjustment of filters via accuracy calculations every 10 trials; ‘Constant’ = Constant formulation of the staircase, where all analysed trials are collected at the same filter number; and ‘Roving’ = Roving staircase, where all trials (across different filter numbers) are used for data analysis. A) Comparison of task accuracy across the data collection methods, where no difference in accuracy was observed between any of the methods (all p > 0.05). B) Comparison of the additional trials required during data collection as percentage of the analysed threshold trials (N.B., no extra trials are collected for the roving staircase design). In Panel B, both the manual and constant staircase methods produced a significant percentage of additional trials (above zero, both p < 0.001), and the constant staircase significantly reduced the number / percentage of these trials compared to the manual adjustment method (both p < 0.001). Black dots are individual data points, while grey areas represent the distribution of values. Large circles denote the group mean in each condition.

Finally, while using a roving staircase design removes the possibility that any additional (non-analyzed) trials will be required, a tighter target accuracy band could be employed to better control the variance in performance accuracy (observed in Figure 7), and the algorithm can be programmed to continue throughout all trials to better control task performance. Therefore, we re-ran the task simulations using a difference in beta cumulative distribution functions between 75-70% (reduced from 80-65%), with a lower bound on the acceptable probabilities increased from 20% to 30%. The results of these simulations compared to the original thresholds can be seen in Supplementary Figure 10, and these changes reduced the simulated standard deviation in task performance accuracy from 8.0 to 6.7.

## Discussion

### Main findings

In this manuscript we demonstrate the utility of pairing a breathing perception task – the ‘Filter detection task’ – with the HMeta-d model of metacognition, to quantify the interoceptive domains of sensitivity, decision bias, metacognitive bias and metacognitive performance. Using simulations, we have shown how the use of a hierarchical model formulation (HMeta-d^18^) can help overcome the challenge of low trial numbers when calculating metacognitive performance metrics using frameworks such as the meta-d model^16^. We demonstrated how this hierarchical model can be extended to include a simultaneous hierarchical estimation of regression parameters linking metacognitive performance to individual difference variables. We also demonstrate the use of the model and appropriate statistics to answer research questions in an empirical dataset of healthy controls and individuals with asthma. However, we did not demonstrate any group differences in our FDT measures (interoceptive sensitivity, interoceptive bias, metacognitive bias or metacognitive performance) within our empirical dataset (including usable data from 56 individuals with asthma and 30 healthy controls), who each completed ∼40 trials (mean ± sd = 42 ± 10) at their perceptual threshold using a manual staircase procedure. Whilst there may be no differences that exist between these groups in the measures tested, the noise associated with using only 40 trials may also mask any underlying differences. Therefore, lastly, we introduce a task algorithm to help target performance accuracy towards 70-75% correct. This accuracy band is optimal for metacognitive analysis as it provides sufficient errors for analysis of confidence-accuracy relationships, while maintaining above-chance performance. We demonstrate the effectiveness of this algorithm using both simulations and via empirical data comparisons when using either manual adjustment strategies or the staircase options provided by the toolbox (constant or roving staircase).

### Computational models of breathing-related interoception and metacognition

As interest in interoception-related research grows across neuroscience, psychiatry, physiology and other scientific communities, the importance of developing robust methodologies for quantification of interoceptive dimensions is paramount. While discussions regarding the validity of tasks such as heartbeat counting in the cardiac domain highlight the need for robust measures of interoception^59^, the FDT offers one route to overcoming some of these issues within the domain of respiration. Here, we highlight the feasibility of applying signal detection theory-derived computational models of both task and metacognitive performance, first introduced by Maniscalco and Lau^16^ (the meta-d model) and derived from theories of ‘Type 2’ performance (distinguishing between one’s own correct and incorrect decisions^60^). Utilizing these signal detection theory models firstly allows us to separate interoceptive sensitivity from decision biases within task (or ‘Type 1’) performance, both of which may be highly informative in disentangling drivers of altered interoception and have been previously quantified using inspiratory loading tasks in controls^61–63^ and children with asthma^64^. Additionally, while there have been reports of possible blunted sensitivity to inspiratory resistive loads with anxiety disorders^65^, there is also an established prevalence of reporting medically unexplained symptoms with anxiety^42^, and even early evidence for a potential relationship between symptom over-report and reduced interoceptive accuracy in healthy individuals^31^. Therefore, as a criterion shift may manifest as differences in interoceptive sensitivity, it is imperative to separate these measures both in healthy individuals and within clinical populations.

While perceptual sensitivity and bias metrics can be directly calculated from behavioural data, the estimation of metacognitive parameters such as meta-d’ often require optimizing a model’s predicted responses to match those observed within the data^16^. However, here we demonstrate that the original meta-d model formulation (using maximum likelihood parameter estimates) is not able to significantly recover group Mratio values below 1 or reliable estimates of individual subject scores when using the low number of trials that are practically feasible within interoceptive experiments (Figure 2). Due to these constraints, here we instead explore the utility of hierarchical formulations of the meta-d model (HMeta-d) derived by Fleming^18^, which can achieve good recovery of metacognitive performance parameters (e.g. Mratio) using as few as 20 trials per subject (Figure 2). Importantly, the meta-d model allows us to differentiate *relative* metacognitive performance (i.e. metacognitive efficiency controlling for task performance) from *absolute* measures of metacognition, such as that calculated from the area under a type 2 ROC curve^15,19,20^. This is important because it is well-established that absolute measures of metacognition may be biased by differences in underlying task performance between individuals or conditions^16^.

Beyond estimating group metrics of metacognition, often it may be desirable to estimate the relationship between individual metacognitive performance and an external measure of interest, for example a clinical score or other behavioural variable. While post-hoc regressions on single-subject parameter estimates are possible, hierarchical models tend to shrink single-subject estimates towards the group mean, thus losing information regarding individual differences and reducing the power of these types of analyses. To this end, we have developed and tested a hierarchical regression model, whereby multiple regression parameters can be simultaneously fit alongside the group logMratio within the HMeta-d model (referred to throughout as RHMeta-d). We find that the sensitivity of the regression model in being able to accurately recover simulated beta coefficients is greatly enhanced when increasing from 20 to 40 and 60 trials, with the width of the posterior (represented by the HDI) notably reducing when trial number is increased (Figure 3). We have also demonstrated the use of this regression approach in an empirical dataset in which interoceptive metacognitive performance was compared against breathlessness symptom reports (measured via the D12 questionnaire) in individuals with asthma (Figure 4).

### FDT toolbox

To aid the use of computational models within interoceptive experiments, we have developed a toolbox to run the FDT according to an accuracy-targeted performance algorithm (freely available for download: https://github.com/ofaull/FDT). While practicalities regarding the step-size of each of the inspiratory resistance filters prevents us from utilizing established psychophysics staircases, we have instead developed an adapted staircase protocol which prompts adjustment of the filter load once the probability falls too far beyond our desired range of 70-75%. As task performance control is carried out online at every trial, any variations in breathing physiology that may alter performance are dynamically accounted for across the task. Both simulations and empirical data show that this algorithm produces performances within the desired range required for employing the computational models described above, where participants need to be performing above chance but below 100% accuracy.

The demonstration of the FDT in the current manuscript utilized a Yes/No task formulation, where a participant is required to answer whether or not a resistance was added to the system (‘Yes’) or stayed the same (‘No’). However, the toolbox also provides the option to employ a two-interval forced choice (2IFC) alternative if desired. While criticisms exist of the application of equal-variance signal detection theory metrics in Yes/No tasks (discussed previously^45^), we also see potential practical utility in using these task variants, for example to quantify measures akin to symptom over- or under-report by estimating the criterion parameter. However, if the metrics calculated from the FDT are to be compared with other perceptual tasks that are run as a 2-interval/alternative forced choice, then the 2IFC option may be desirable, allowing for comparable model assumptions across tasks.

Lastly, the toolbox also offers two alternative staircase options to control task performance in either a Yes/No or 2IFC formulation. In the original protocol presented by Garfinkel and colleagues^15^, the aim was to collect 20 usable trials at a specific number of filters where performance first fell below 75%, thus corresponding to the participant’s perceptual threshold and providing a measure of interoceptive sensitivity. Whilst the number of additional (unused) trials required can be greatly reduced by employing the adapted staircase algorithm in the toolbox (Figure 7), an alternative approach is to employ a ‘roving’ staircase, whereby all trials are used in the calculation of interoceptive measures, and interoceptive sensitivity is taken as an average of the filter numbers used across trials. As the risk of needing additional trials is removed, this approach allows experimenters to tighten accuracy thresholds to improve task performance control, as the aim of this staircase is no longer to find a single filter that elicits the desired accuracy. This roving staircase option would likely prove a more viable alternative if using the FDT in a clinical setting, removing the possibility of any additional trials while maintaining adequate representations of interoceptive sensitivity. We note however that roving staircases also have potential downsides in artificially inflating estimates of metacognitive sensitivity when compared to constant-stimulus designs (see Rahnev and Fleming^66^ for further discussion of this issue).

### Limitations

While this experimental setup provides a progression towards measuring quantities related to interoception of breathing, a number of limitations exist that could be addressed in future work. The first of these is that while the resistance applied is static, the resulting pressure differential across the resistance is flow-dependent, such that larger inspiratory flow will generate larger inspiratory pressure differences (see Supplementary Material for further details). Furthermore, inherent resting resistance and inspiratory pressures are also variable between individuals, depending on factors such as anatomical structure of the airways and physiological differences in inspiratory musculature. Therefore, if measures of inspiratory pressure and flow were added to the system, more detailed quantification of perceptual sensitivity may be determined by considering the changes in both the inspiratory pressure and flow (relative to the baseline breaths) that were required to detect the resistance. The use of mouth pressure, in particular, could be used as a replacement for the number of filters as a more nuanced measure of interoceptive sensitivity. As pressure will change in response to both changes in resistance as well as inspiratory flow (see Supplementary Figure 4 for details), this measure would thus incorporate any inter-filter and inter-participant inspiratory flow variability. However, it is worth noting that the addition of these physiological measures would not alter the metacognitive performance scores, as both controlling task performance and accounting for any remaining performance variability (by using the meta-d model) renders these independent of interoceptive sensitivity^17^. The downside of these additional measures would be the loss of some of the task simplicity, and thus its feasibility for use in a wide range of settings. A further notable limitation of the current version of the FDT is the time required for completion of the task.

The original manual staircase proposed by Garfinkel and colleagues^15^ (used in the collection of the asthma data presented here) could require more than 60 minutes to acquire the final 20 trials used in the analysis, and other inspiratory load paradigms measuring only objective performance also require up to 60 minutes to collect only 7 trials at each level of resistance^12^. Using this manual staircase and ∼40 trials in each participant, we were unable to identify any differences in interoceptive sensitivity, interoceptive bias, metacognitive bias or metacognitive performance between individuals with asthma and healthy controls. Additionally, while worsened D12 scores appeared to relate to reduced metacognitive performance in asthma (Figure 5), this relationship did not reach statistical significance. To aid the collection of more trials in order to reduce the noise associated with each of our measures, the implementation of the FDT staircase allows for up to 60 usable trials in approximately the same amount of time (45-70 minutes). To further improve the ecological validity of the task, a roving staircase design with only 40 trials could be employed; this would dramatically reduce the time required for task completion to approximately 20-30 minutes per participant.

An additional limitation of the current design is the currently fixed staircase step-sizes induced by adding or removing a filter, and their lack of highly-accurate factory calibrations. While the resistance increments of 0.22 cm H_2_O / L.sec^-1^ provided here (see Supplementary Material for details) are smaller than those used in previous paradigms such as that by Davenport et al. (0.20 – 11.46 cm H_2_O / L.sec^-1^)^12^, sophisticated electronic devices that can deliver very variable small resistances (<0.5 cm H_2_O / L.sec^-1^) are not yet widely available. The development of such a device would allow for variable step-sizes, reduce the time required to manually change filters, and allow greater control over inspiratory flow rate and volume to reduce breath-by-breath variability. Such devices may either incrementally change resistance using techniques such as an adjustable aperture, or with even further sophistication establish a constant inspiratory resistance via biofeedback devices. The possible improvements in control over the staircase step-size may also allow for more established staircase procedures (such as QUEST) to be implemented here, where the fine-grain coverage of perceptual evidence strength required to accurately identify perceptual threshold may then become available. Furthermore, automated developments would also greatly reduce the current experimenter burden of having to manually change filters and interact with the task algorithm at every trial.

While the limitations in the current measures of perceptual sensitivity are worth observing and improving, these limitations do not discount the utility of the current measures. While the noise of the perceptual sensitivity metrics will be notable (but not necessarily insurmountable), the control of task performance allows the metacognitive measures to be somewhat independent of this noise. Furthermore, the measure of metacognitive performance directly accounts for any remaining differences in task performance by creating a ratio of meta-d’ / d’. Finally, keeping participants comfortable and reinforcing the notion that the task should be performed with normal pace and depth of breathing should limit large differences in inspiratory flow and pressure, and filters could be additionally numbered to ensure consistency in incremental steps.

## Conclusions

Here we present a breathing-related interoceptive application of a computational model designed to tease apart important aspects of perception: sensitivity, decision bias, metacognitive bias and metacognitive performance. Whilst interoceptive experiments often suffer from low trial numbers, by combining a breathing perception task with a hierarchical statistical model we were able to develop a robust algorithm to control task performance while maximizing the number of useful trials for analysis.

The FDT toolbox is freely available for download (https://github.com/ofaull/FDT), as are the statistical methods employed (MLE model: http://www.columbia.edu/~bsm2105/type2sdt/; HMeta-d and RHMeta-d: https://github.com/metacoglab/HMeta-d/).

## Acknowledgements

Data provided in this manuscript is not yet publicly available: Please contact authors for data access. OKH (née Faull) was supported as a Marie Skłodowska-Curie Postdoctoral Fellow from the European Union’s Horizon 2020 research and innovation programme (under the Grant Agreement No 793580), and as a Rutherford Discovery Postdoctoral Fellow from the Royal Society of New Zealand. MA is supported by a Lundbeckfonden Fellowship (R272-2017-4345), and an AIAS-COFUND II fellowship, which is supported by the Marie Skłodowska-Curie actions under the European Union’s Horizon 2020 (Grant agreement no 754513), and the Aarhus University Research Foundation. SJH was supported by the grant #2017-403 of the Strategic Focal Area “Personalized Health and Related Technologies (PHRT)” of the ETH Domain. KES is supported by the René and Susanne Braginsky Foundation and the University of Zurich. KTSP was supported by the JABBS Foundation for this work. SMF is supported by a Sir Henry Dale Fellowship jointly funded by the Wellcome Trust and Royal Society (206648/Z/17/Z).

## Supplementary Material

**Supplementary Figure 1.**
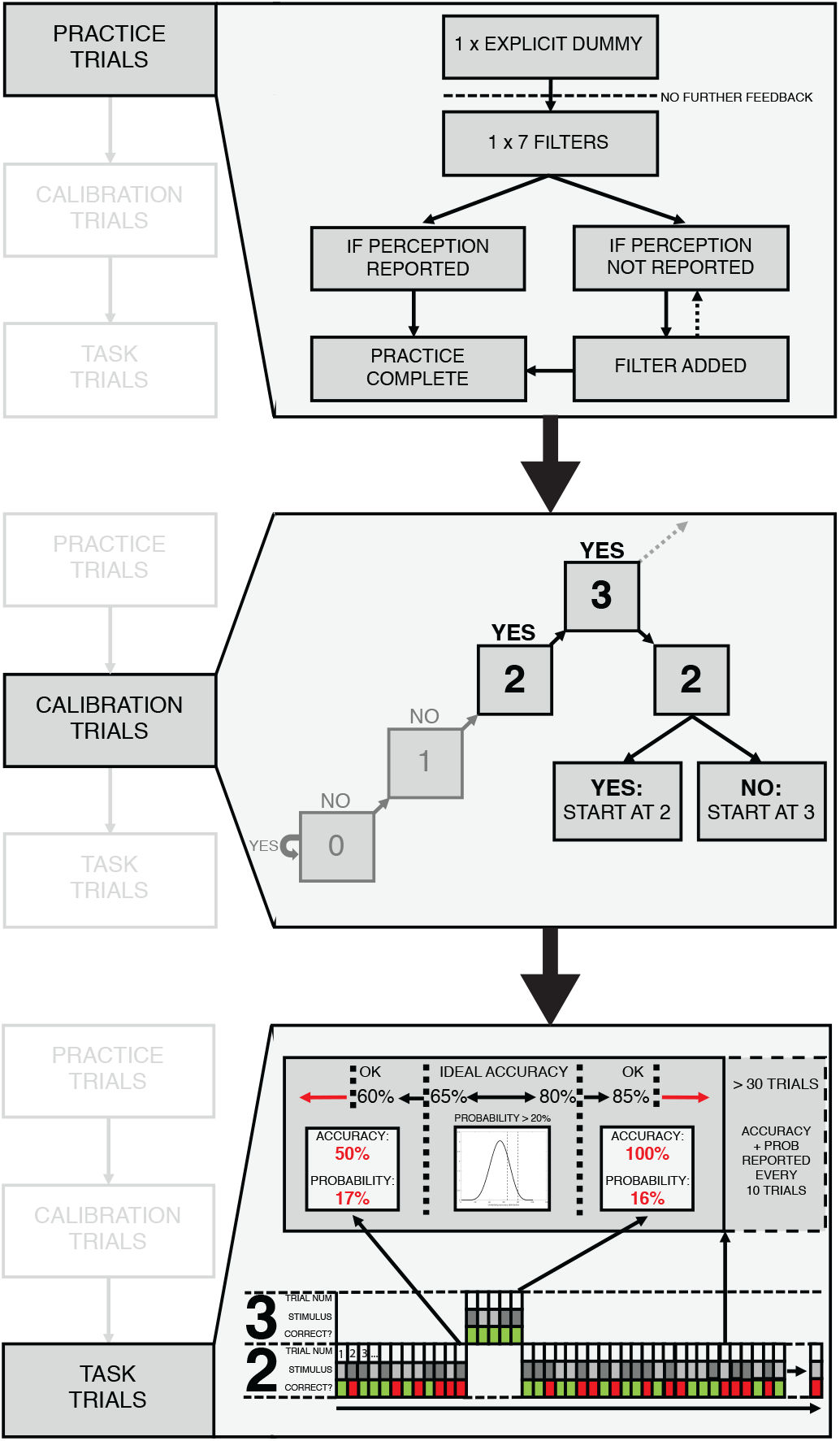
Visualization of the task performance algorithm. TOP: Practice trials, consisting of an explicit dummy (where the participants are told a sham resistance is added), followed by a large load (7 filters) where no feedback is given (this is then maintained for the rest of the experiment). If no resistance is perceived in the Yes/No format or the answer is incorrect in the 2IFC format, filters are added until a correct resistance is reported. MIDDLE: Calibration trials, where (starting from the dummy), filters are added until two consecutive resistances are reported (i.e. two ‘Yes’ answers for the Yes/No format of the task). If performing a 2IFC task, the required ‘Yes’ answers for the calibration trials are replaced by correct 2IFC answers. Following this, one final calibration trial is performed with one less filter, to determine the starting value for the task trials. BOTTOM: Task trials, where cumulative task accuracy at each trial is transformed (using a beta distribution) into the distribution of underlying accuracies that could have produced the task performance. An upper bound (here 80%) and a lower bound (here 60%) is used to calculate the probability that the participant is performing at the targeted accuracy. If this probability falls below the error risk threshold (here 20%), a filter change is prompted – either the addition of a filter if the accuracy is too low, or the removal of a filter if the accuracy is too high. This continues until either a specified number of trials (here 60 trials) are completed at either one filter number (using the ‘constant staircase’ task design) or at a range of filter numbers (using the ‘roving staircase’ task design), the latter requiring no additional trials to be measured that will not be used in the analysis. If a constant staircase is used, the algorithm is stopped at 30 trials, and experimenter intervention can occur every 10 trials subsequently if task performance is drastically altered and no longer deemed acceptable.

### Filter resistance notes

Detailed information describing the spirometry filters used for this task is provided by GVS (http://www.gvs.com/images/uploads/user/Catalogues%20Pdf/Cat%202019%20Healthcare%20Air.pdf; product number 2800/22BAUF). A copy of the relevant information is additionally outlined in Supplementary Figure 2 (below) for reference.

As resistance is the force that opposes the movement of a fluid (or particles) and is a mechanical property of the circuit, this remains constant over the course of each trial. However, the inspiratory force exerted in each breath can change inspiratory pressure, which leads to changes in inspiratory flow (as determined by the circuit’s resistance). The relationship between flow (*Q*), differential pressure (*P*) and resistance (*R*) is determined by Ohm’s law:

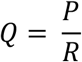

Utilizing this relationship, we can use the reference values provided in Supplementary Figure 2 to calculate the resistance for each filter as < 0.48 cm H_2_O / L.sec^-1^ at 30 L.min^-1^ (0.5 L.sec^-1^), calculated as *R* = *P* ÷ *Q*.

**Supplementary Figure 2.**
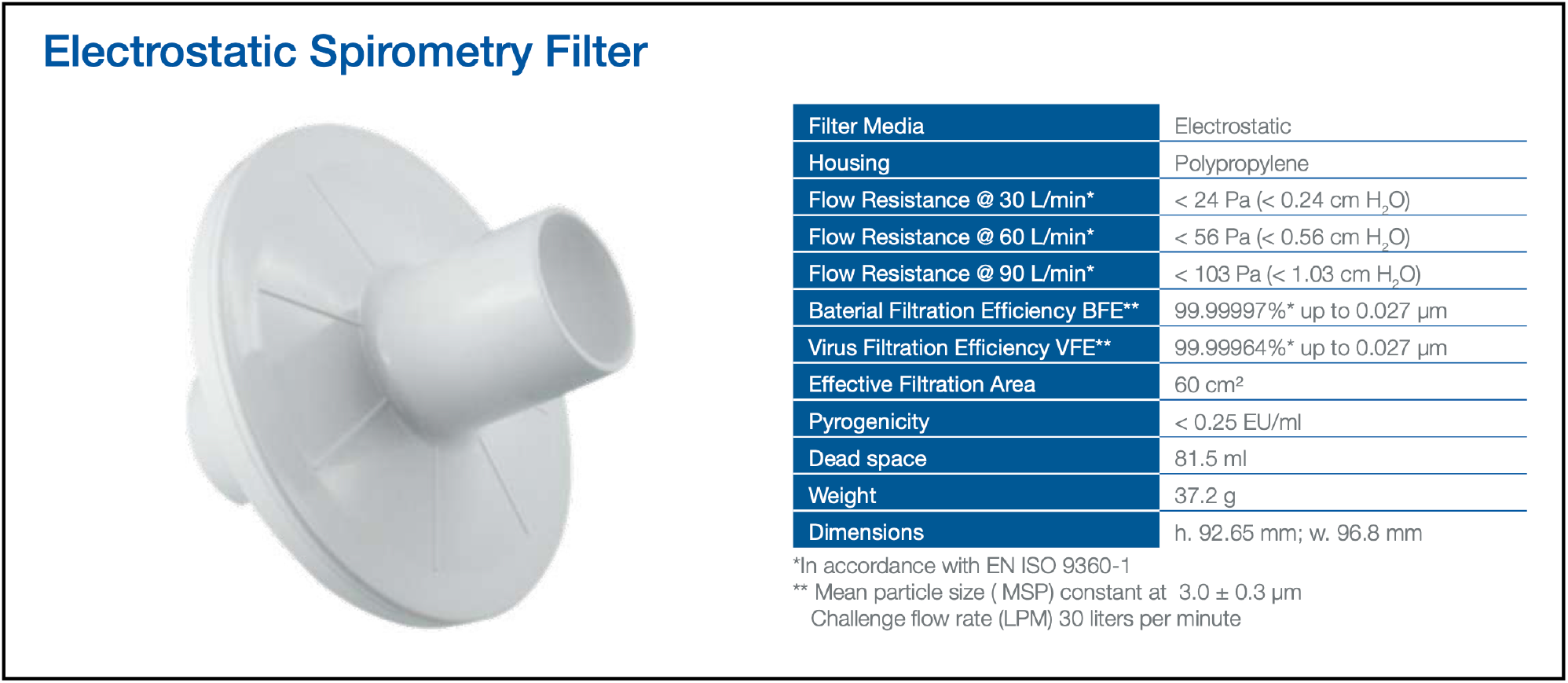
Details of the spirometer filter product used to add resistance for detection in the FDT. Taken from GVS Filter Technology product guide, and details can additionally be found listed on the webpage (http://www.gvs.com/product-family/187/679/2800). The “Flow Resistance” measures provided in this table are pressure values, and thus resistance can be calculated by dividing the given pressure values by the given flow rate.

Additionally, we measured the inspiratory pressure and flow across a range of filters over all levels, such that we were able to measure and calculate inspiratory resistance values. Resistance values are presented in Supplementary Figure 3A, and we demonstrate that each additional filter provides an added resistance of 0.22 cm H_2_O / L.sec^-1^, with the baseline resistance of the circuit (including the dummy filter) 0.31 cm H_2_O / L.sec^-1^. Furthermore, Supplementary Figure 3B shows clear separation between the resistance values calculated for each filter level, although measurement error from pressure and flow devices contributes to a small amount of variability in the resistance magnitude.

Finally, we demonstrate the strong relationship between increases in inspiratory flow and pressure (Supplementary Figure 4) using an example filter level of 5. If inspiratory flow is increased, inspiratory pressure will also increase proportionately against this static resistance. Therefore, the magnitude of inspiratory pressure at perceptual threshold (∼70% performance accuracy) would be a more accurate measure of interoceptive sensitivity, as this would take into account the effect of inspiratory flow as well as the number of filters used for an external resistance.

**Supplementary Figure 3.**
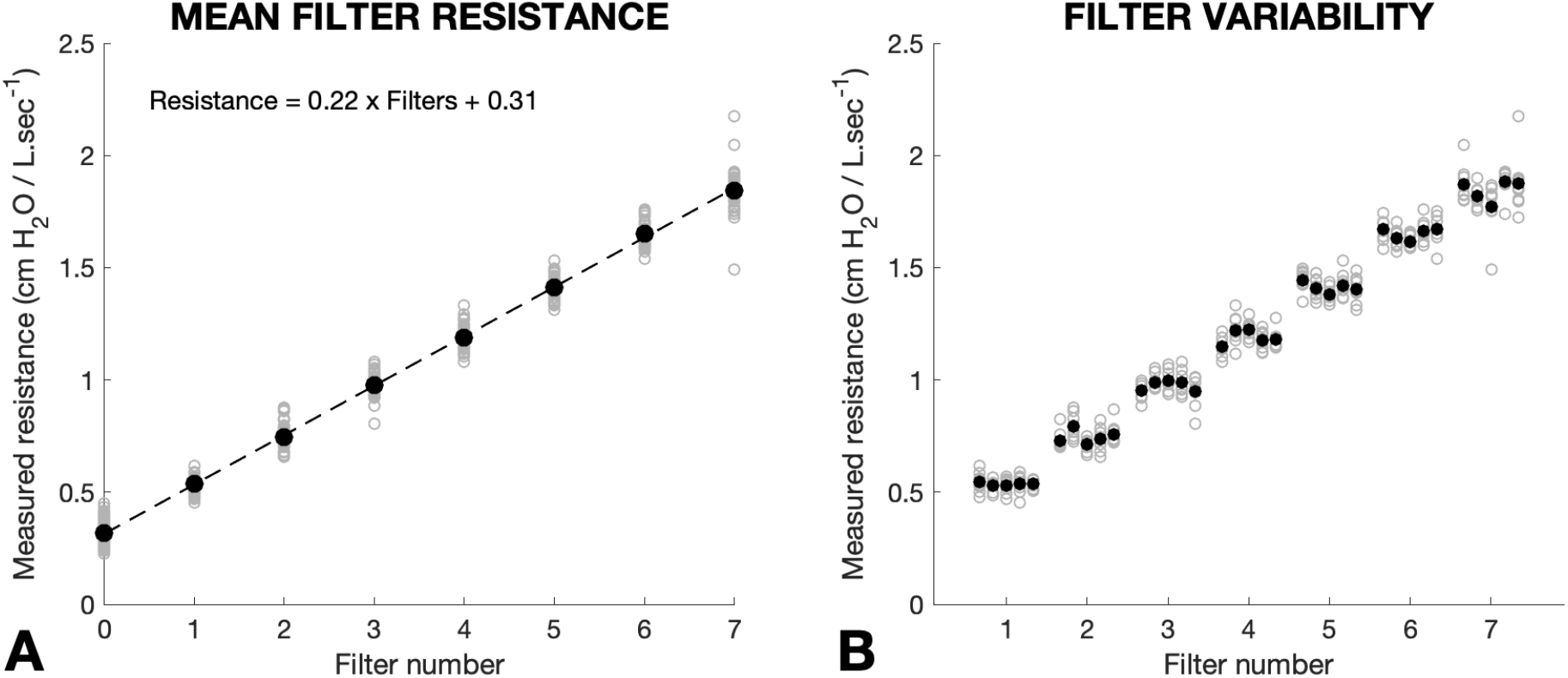
Magnitude of inspiratory resistance induced by increasing filter levels. A) Grey circles indicate the resistance values calculated for each trial at each filter level, with black circles denoting the mean value at each level. The dotted line represents a regression of the filter level against the mean resistance values, such that the slope and intercept (resistance of the circuit with the dummy attached) can be quantified. B) Grey circles indicate the resistance values calculated for each trial using five different filter combinations at each filter level, with black circles denoting the mean value for each filter combination. Clear separation can be seen between the resistance produced at each level of filters.

**Supplementary Figure 4.**
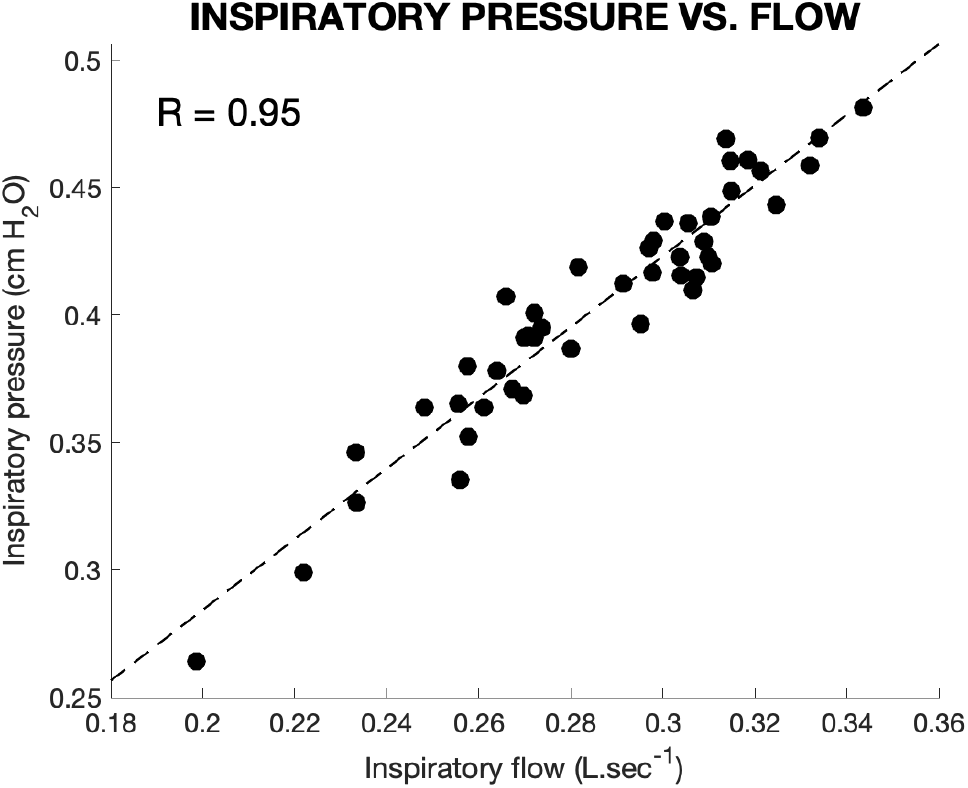
Relationship between inspiratory flow and inspiratory pressure using an example of 5 filters. The inspiratory pressure generated is proportional to the magnitude of inspiratory flow, and thus measures of inspiratory pressure at perceptual threshold would provide a more accurate measure of interoceptive sensitivity than the number of filters used, as this will account for variations in inspiratory flow.

**Supplementary Figure 5.**
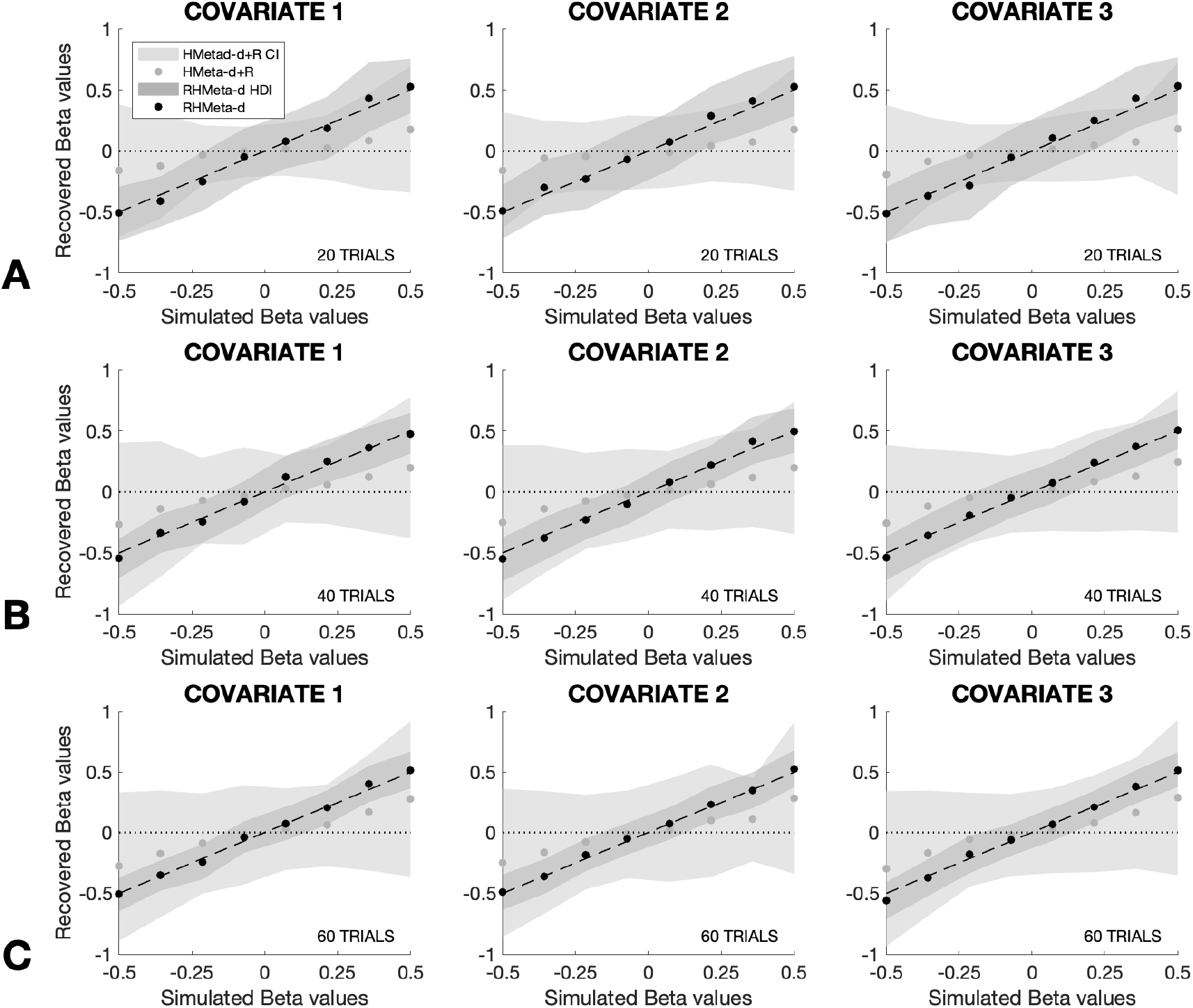
Demonstration of the recovery of three group regression parameters (betas) against the logMratio using either the original HMeta-d model combined with a post-fit linear regression (HMeta-d+R), or an extended multiple regression HMeta-d (RHMeta-d) model, the latter where group linear regression parameters are simultaneously hierarchically fit alongside logMratio. Results are shown for simulations using 20 trials (A), 40 trials (B) and 60 trials (C). Grey areas denote the 95% highest density interval of the sampled estimate. Dashed lines represent ideal recovery of group beta values, and dotted lines at zero demonstrating the ability of the model fit to significantly recover group estimates (with highest density intervals not including zero).

**Supplementary Figure 6.**
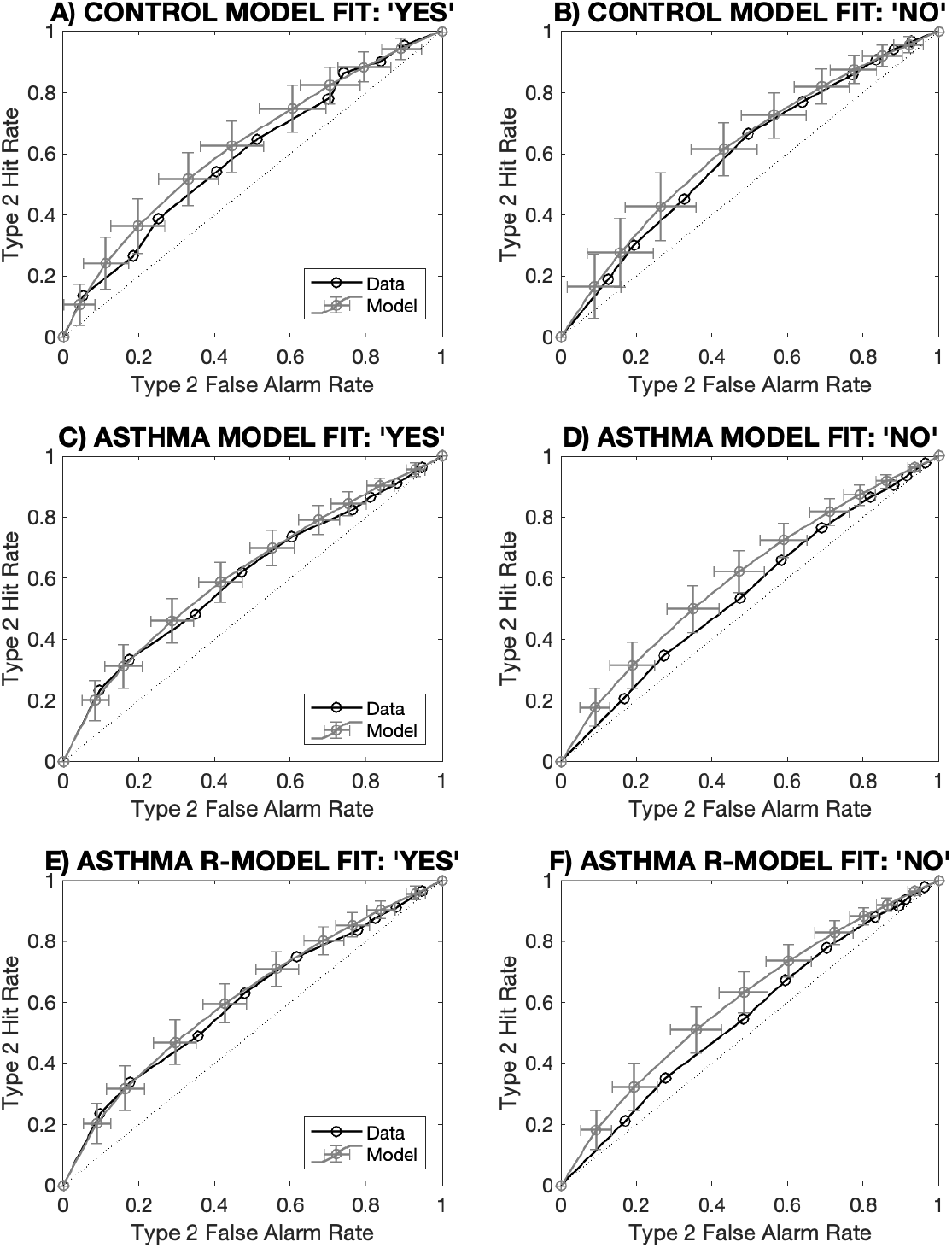
Demonstrations of the model fits by comparing the observed and model estimates of the Type 2 ROC curves for both ‘Yes’ and ‘No’ responses (regarding the presence of an added inspiratory resistance). Model fits are shown for the control data fit (A and B), the asthma data fit (C and D), and for the asthma data extended regression model (E and F).

**Supplementary Figure 7.**
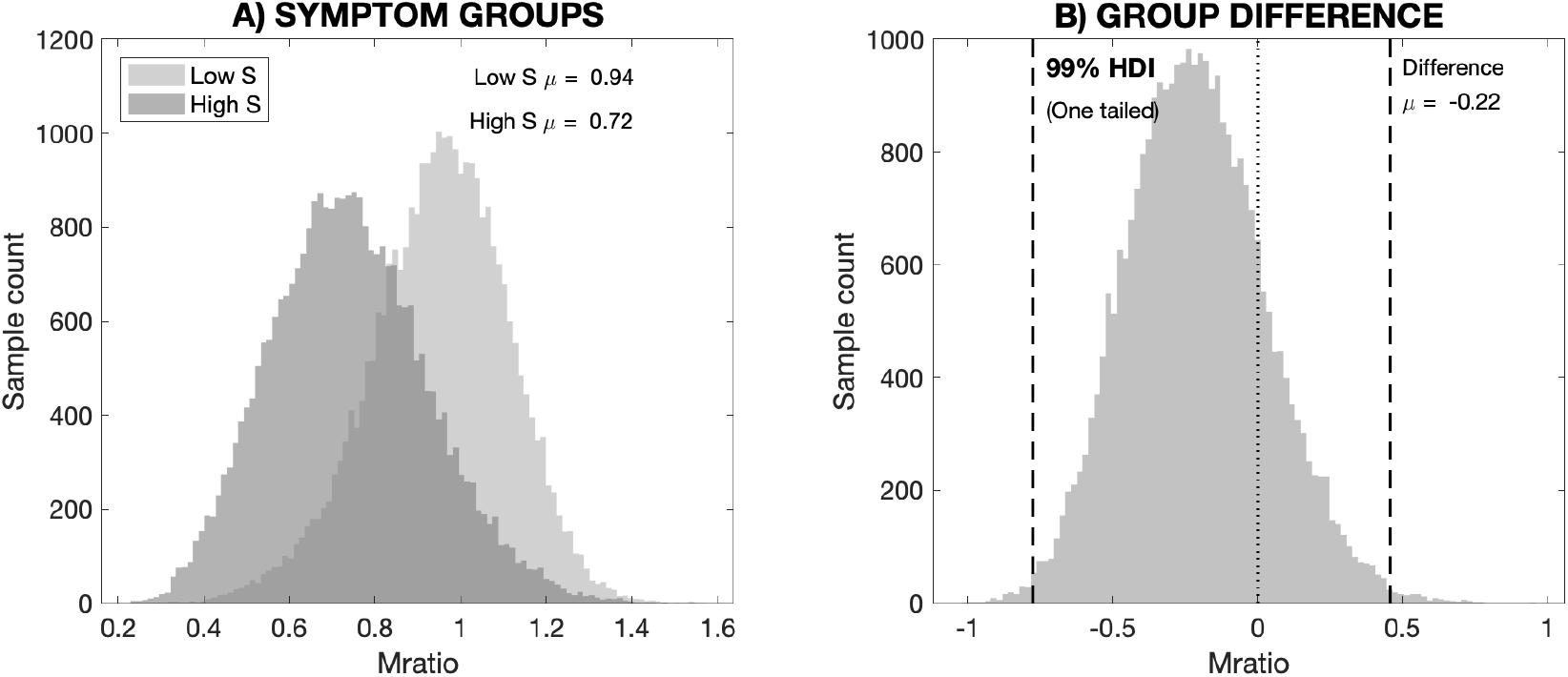
Comparisons between metacognitive performance (logMratio) and symptom load in asthma, using a median-split analysis for high and low symptoms. A) The sampled posteriors of the hierarchically-estimated metacognitive performance parameter (Mratio) for individuals with > median D12 scores (high symptom load group) or < median D12 score (low symptom load group). B) The distribution resulting from the difference between the group Mratio distributions, where the dashed lines represent the two-tailed non-significant 99% highest density interval (HDI), and the dotted line denotes zero.

**Supplementary Figure 8.**
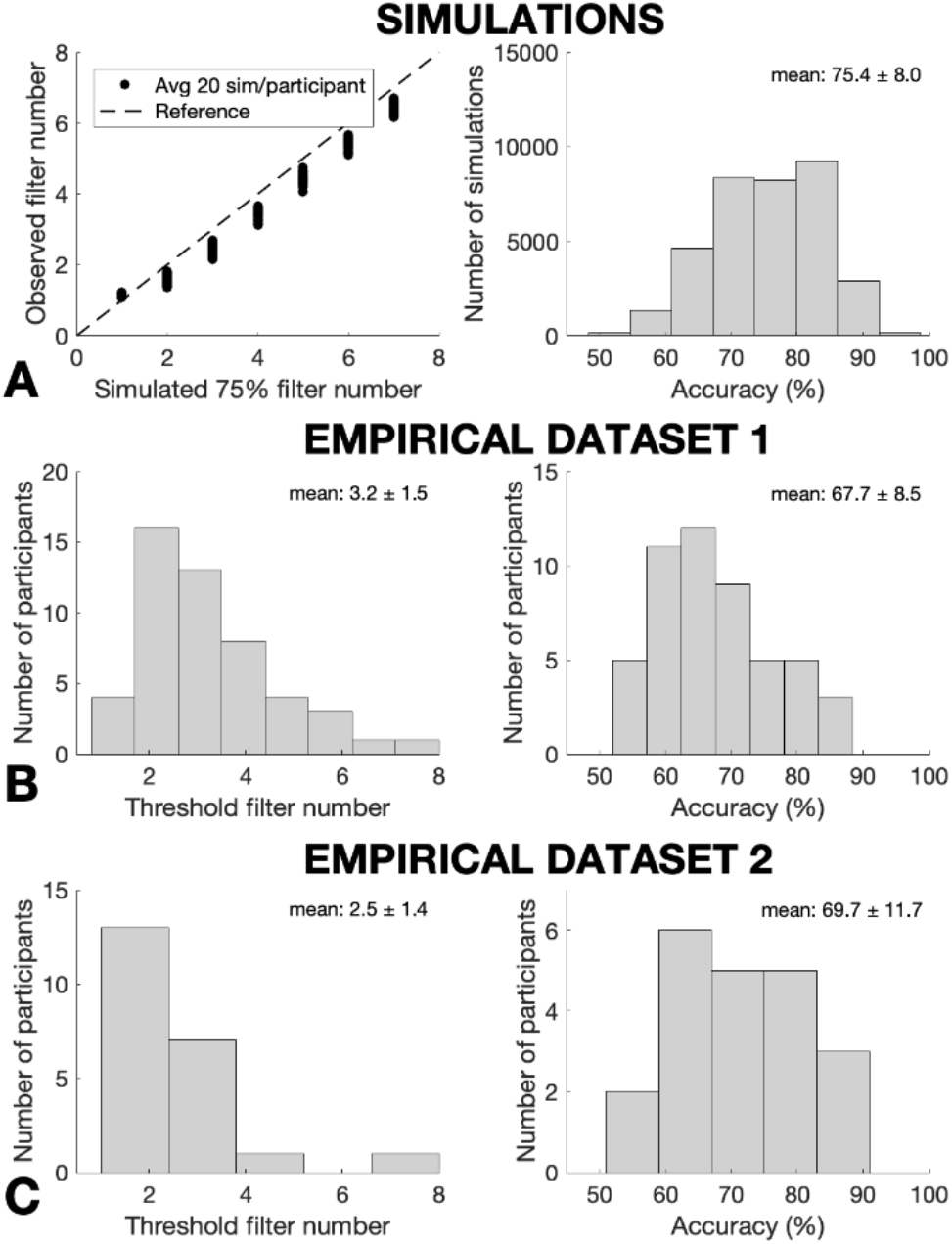
Results demonstrating the use of an adapted roving staircase algorithm for targeted task difficulty over 60 trials. A) Simulation results, where data were generated from a range of logistic sigmoid functions bounded between 0.5 and 1, with 20 simulations for each sigmoid (‘participant’) from each of five starting points – from two filters below to two filters above the 75% threshold filter. Left: Simulated and recovered 75% filter number for each simulated ‘participant’. Right: Histogram of the task accuracy scores for the 60 threshold trials for all simulations. B) Data collected using a Yes/No version of the task (with a constant staircase, but where the first 60 trials – regardless of filter number – were analyzed to represent a roving staircase design), where 50 participants were measured. Left: Histogram of the measured threshold filter number for each participant. Right: Histogram of the task accuracy scores for the 60 threshold trials for the 50 measured participants. C) Data collected using a Yes/No version of the task (with a roving staircase), where 20 participants each completed 60 trials (total). Left: Histogram of the measured threshold filter number for each participant. Right: Histogram of the task accuracy scores for the 60 threshold trials for the 50 measured participants. All histograms are reported with mean ± standard deviation.

**Supplementary Figure 9.**
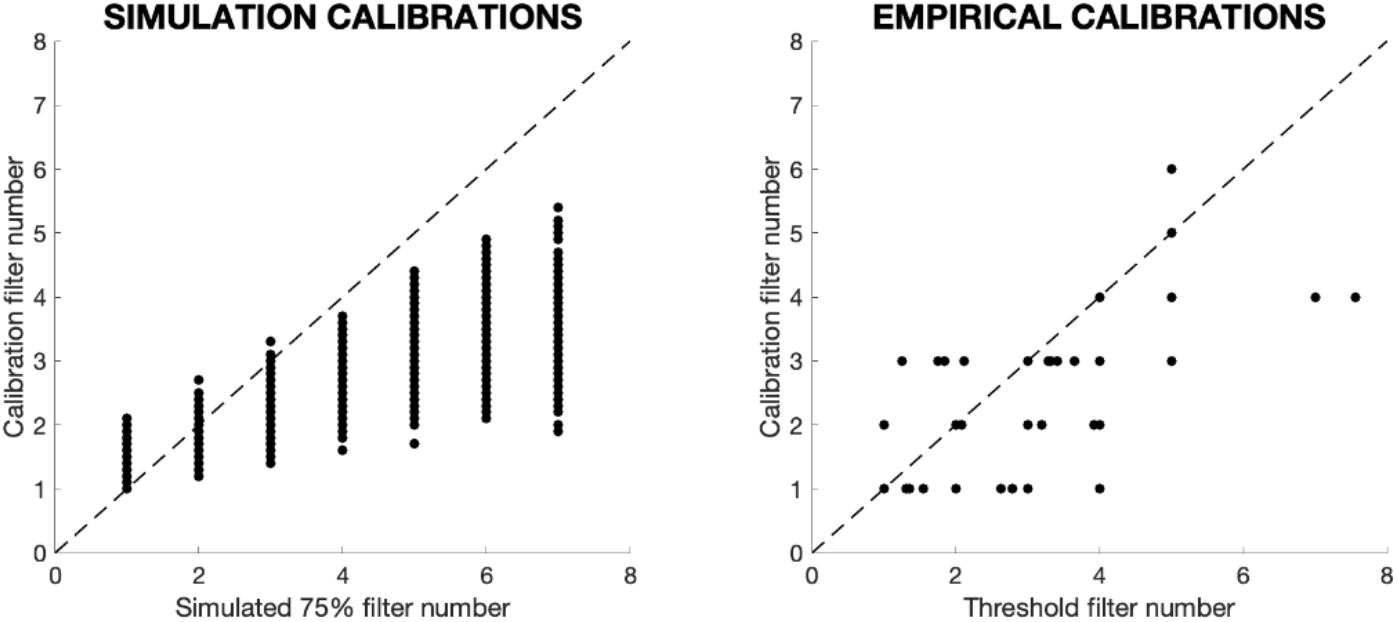
Results for the calibration algorithm from both simulated and empirical data. Left: Simulated calibration filter plotted against the threshold (75%) filter, demonstrating a bias towards lower calibration values. Right: Empirical calibration results plotted against the task threshold filter number for data from both constant and roving staircase designs (roving staircase threshold filter numbers are calculated as the average filter number across the task trials).

**Supplementary Figure 10.**
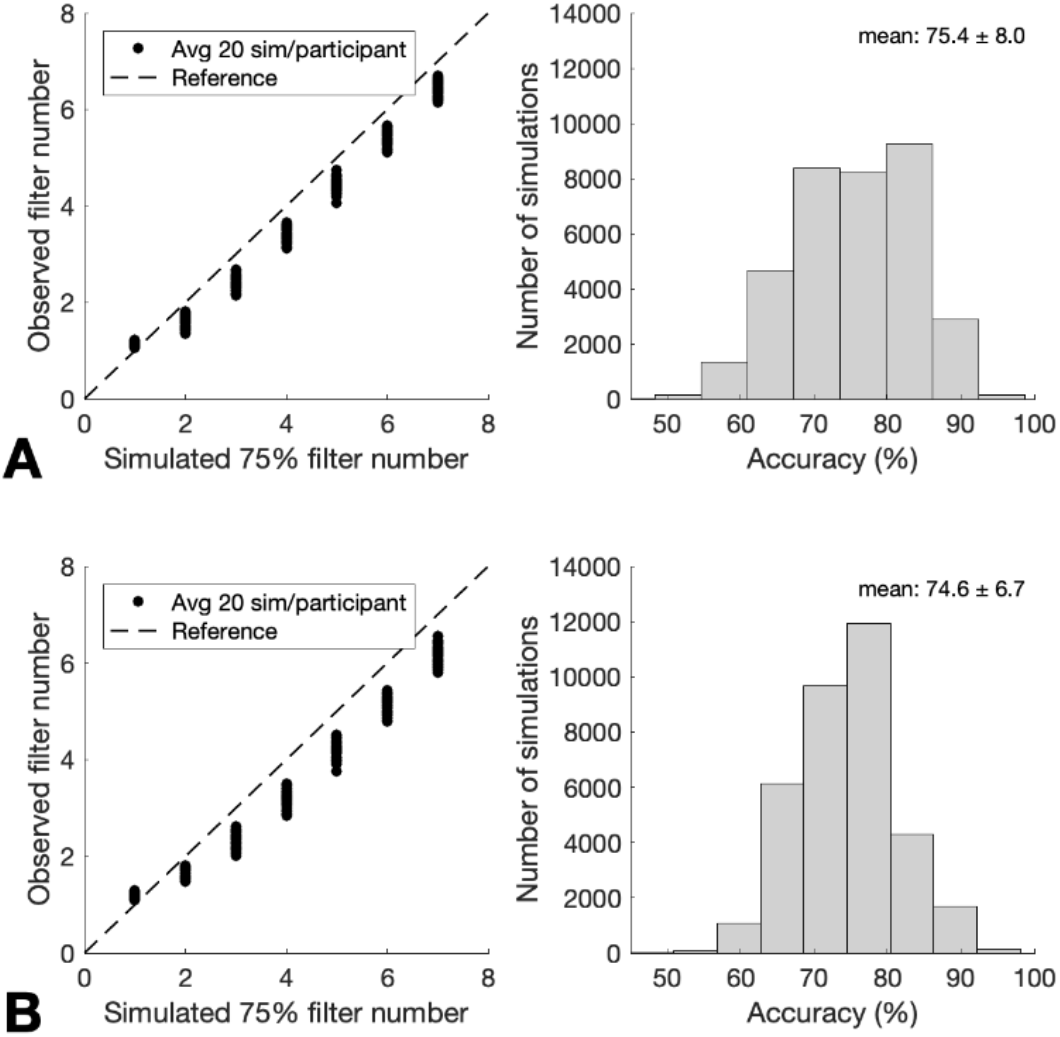
Simulated results demonstrating the use of an adapted roving staircase algorithm for targeted task difficulty over 60 trials, using two different sets of thresholds for prompting a change in filter number. In both panels, data were generated from a range of logistic sigmoid functions bounded between 0.5 and 1, with 20 simulations for each sigmoid (‘participant’) from each of five starting points – from two filters below to two filters above the 75% threshold filter. A) Data were generated using an upper bound of 80% and a lower bound of 65% on the beta distributions calculated from task performance scores, using 20% as a false positive threshold where probabilities below this threshold prompted a filter change. B) Data were generated using an upper bound of 75% and a lower bound of 70% on the beta distributions calculated from task performance scores, using 30% as a false positive threshold where probabilities below this threshold prompted a filter change. Both panels – Left: Simulated and recovered 75% filter number for each simulated ‘participant’. Right: Histogram of the task accuracy scores for the 60 threshold trials for all simulations. Data from B show a very similar mean and reduced standard deviation for the resulting task performance accuracies generated.

